# Environmental selection and epistasis in an empirical phenotype-environment-fitness landscape

**DOI:** 10.1101/2021.04.14.439889

**Authors:** J.Z. Chen, D.M. Fowler, N. Tokuriki

## Abstract

Fitness landscapes, mappings of genotype/phenotype to their effects on fitness, are invaluable concepts in evolutionary biochemistry. Though widely discussed, measurements of phenotype-fitness landscapes in proteins remain scarce. Here, we quantify all single mutational effects on fitness and phenotype of VIM-2 β-lactamase across a 64-fold range of ampicillin concentrations. We then construct a phenotype-fitness landscape that takes variations in environmental selection pressure into account. We found that a simple, empirical landscape accurately models the ~39,000 mutational data points, suggesting the evolution of VIM-2 can be predicted based on the selection environment. Our landscape provides new quantitative knowledge on the evolution of the β-lactamases and proteins in general, particularly their evolutionary dynamics under sub-inhibitory antibiotic concentrations, as well as the mechanisms and environmental dependence of nonspecific epistasis.

**One Sentence Summary:** An empirical fitness landscape discloses the environmental dependence of mutational effects in VIM-2 β-lactamase.

## Main Text

The quantitative understanding of genotype-phenotype-fitness relationships of proteins is central in evolutionary biochemistry (*1*–*3*). Genotype, a protein’s sequence information, largely dictates its phenotypes, which are the measurable *in vitro* physio-chemical traits of proteins such as protein activity and stability. Nonetheless, *in vivo*, multiple factors could modulate protein function and behavior in the cell (*e.g.*, the level of protein expression). The fitness of the organism carrying a specific protein is greatly influenced by the environment where the organism lives, and the selection pressure to a particular protein’s function vary significantly with environmental factors. Thus, deciphering the ‘phenotype-fitness’ relationship across environmental variation is essential to understanding and potentially predicting a protein’s evolutionary dynamics (*4*–*8*). The phenotype-fitness landscapes typically manifest as a non-linear relationship, in the form of a ‘threshold’ or ‘mesa-like’ model (*6*, *9*–*14*). In this model, the fitness conferred by variants with biochemical traits above a certain selection ‘threshold’ would saturate or plateau, while fitness rapidly decreases for variants below the threshold of the trait. Such models have been widely recognized conceptually, and provide molecular explanations for important evolutionary phenomenon such as nonspecific epistasis (*4*, *14*–*18*), mutational robustness (*9*, *11*, *12*, *15*, *18*, *19*) and evolvability (*6*, *10*, *12*, *13*, *20*–*22*). However, quantitative and experimental studies of the threshold model are rare, as most reports contain just a handful of data points and each landscape is confined to the scope of a single level of selection pressure (*9*, *10*, *16*, *19*, *22*). Consequently, detailed empirical models that can recapitulate the fitness landscapes of proteins, let alone the phenotype-fitness relationship across varying environmental conditions, have never been established. Such models will enable the quantitative evaluation of the evolutionary dynamics in proteins, and the development of predictive models for protein evolution in the response to a given environmental selection pressure.

To describe a realistic phenotype-fitness landscape, we require a quantitative mapping of the phenotype-fitness relationship with careful examinations in the context of environmental factors. Recent advances in largescale mutational characterization experiments, such as deep mutational scanning (DMS)(*23*, *24*), have enabled the high-throughput measurement of protein fitness (growth of host organism carrying encoded protein variants) and phenotype (*e.g.*, antibiotic resistance level, protein binding and expression) for several thousands of mutational variants (*16*, *20*, *25*–*34*). DMS across different conditions or genetic backgrounds can also determine higher dimensional characteristics of protein phenotype and/or fitness, such as antibiotic concentration dependent fitness(*20*, *25*, *32*), substrate specificity(*25*, *35*, *36*) and inter/intra-protein and protein-environment interactions (*17*, *27*, *28*, *37*–*43*).

In this work, we seek to obtain a global phenotype-environment-fitness landscape using the VIM-2 β-lactamase as a model system, for which we previously generated all single point mutants (~5600 variants) (*35*). To this end, we determined the fitness of an *Escherichia coli* strain carrying VIM-2 variants across different environments dictated by seven different ampicillin (AMP) concentrations (~39,000 data points). We also measured the phenotype, specifically the antibiotic resistance level (*EC*_50_), of each variant by integrating the experimental data across all tested AMP concentrations into dose-response curves. We show that the phenotype-fitness relationships exhibit a universal behavior according to AMP concentration. Taking advantage of this result, we generate an empirical model for generalizing the phenotype-environment-fitness landscape of VIM-2. We then demonstrate that this empirical and quantitative model can explain the evolutionary behavior of the resistance gene at low antibiotic concentrations, and the molecular mechanisms underlying the prevalence of nonspecific epistasis in mutational datasets.

## Results

### Selection of VIM-2 variant libraries in a range of ampicillin concentrations

The overall workflow for our antibiotic selection experiment is presented in **Fig. 1A**. We previously generated a library containing ~5600 single amino acid variants of wild-type (*wt*) VIM-2 in a vector that expresses VIM-2 through a constitutive ampR promoter (*35*). The library was transformed into *E. coli* and subsequently grown in LB liquid media containing a range of AMP concentrations (4.0-256 μg/mL AMP in two-fold increments), or in the absence of AMP, for 6 hours at 37°C. The AMP concentrations used in this study covers the full dynamic range of VIM-2’s conferred resistance, from just above 2 μg/mL AMP where *E. coli* can grow without VIM-2 expression (*35*), to more than double the *EC*_50_ of 84 μg/mL where *E. coli* expressing *wt*VIM-2 can no longer grow. The concentration of cells in each culture after the selection was measured via *OD*_600_ to track the total population size (**Fig. 1B**), and the mutated region of VIM-2 gene was amplified by PCR for deep sequencing (*35*). Two replicates of the experiment with antibiotic selection and three replicates of growth without selection were conducted.

**Fig. 1.**
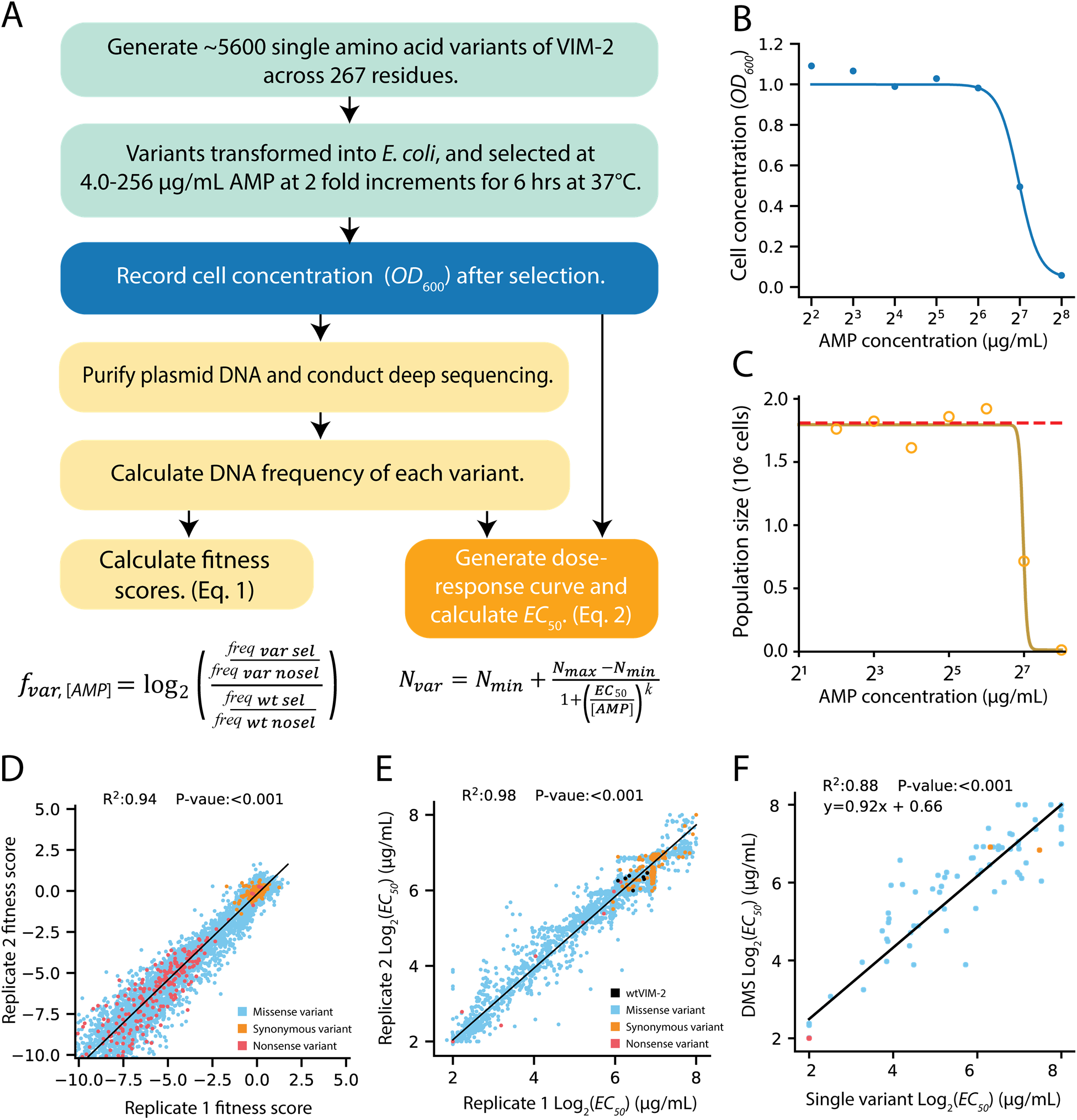
Workflow for generating fitness scores and *EC*_50_ values using DMS. (**A**) Flowchart of the DMS process. A library of protein variants is first generated and transformed into host *E. coli*, which are then placed under selection with a range of antibiotic concentrations. The *OD*_600_ of the selected culture is recorded after selection and the selected culture is processed for deep sequencing. For each variant, the DNA frequency can be used to directly calculate the fitness scores, as well as the population size of each variant (*N_var_*) after selection (using the *OD*_600_). The population size can be used to generate a dose-response curve for each variant. (**B**) Cell density of a culture pool encompassing variants in the first 39 positions of VIM-2, measured after selection for each AMP concentration. (**C**) Example of a dose-response curve generated for the I19A variant. The solid line represents a fit to a sigmoidal curve. The red dashed line indicates the average non-selected population size of the replicate. (**D**) Correlation of variant fitness scores between two replicates under selection with 64 μg/mL AMP. For panels D-F, variants are colored by mutation type; the same color scheme applies to all panels. (**E**) Correlation of *EC*_50_ values fitted from two replicates. (**F**) Correlation between *EC*_50_ determined by using deep sequencing (y-axis) and or from isolated variants grown in separate liquid cultures (x-axis).

### Calculation of fitness scores and EC_50_

The output of the DMS experiments generates sequencing counts for each variant in the library, which can be used to calculate the fitness scores (reflecting relative growth) at each AMP concentration as well as the phenotype (*EC*_50_) for each variant. The crux of our approach is that the *EC*_50_ values are calculated independently from the fitness score in each condition, such that their values are not interdependent, even when using the same set of deep sequencing data. Thus, this allows us to examine the relationship between phenotype and fitness as separate factors under each selection pressure.

The fitness (*f*) of a variant under each AMP concentration is calculated as the relative growth of a variant compared to *wt*VIM-2 as we described previously (**Fig. 1A**, **Data S1** for all fitness scores) (*35*). For each variant, and at each level of AMP selection, an enrichment score was calculated as the ratio of the frequency of a variant after selection to the frequency without selection. Next, the fitness score of each variant was calculated by normalizing the enrichment score of a variant to that of *wt*VIM-2, generating a measure of fitness relative to *wt*VIM-2 using **equation (1)**.

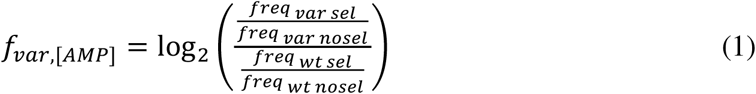

The antibiotic resistance level of each variant was determined via a dose-response curve by integrating the deep sequencing data from all seven different conditions (**Fig. 1A-1C, Fig S1**, **Data S2** for all *EC*_50_ values). To this end, we first calculated the population size of cells expressing each variant (*N*_var_) at each antibiotic concentration by multiplying the frequency with the cell concentration of the culture after antibiotic selection. Then we generated a dose-response curve for each variant by plotting *N*_var_ against the antibiotic concentration, and calculated the *EC*_50_ by fitting the sigmoidal relationship in **equation (2)**

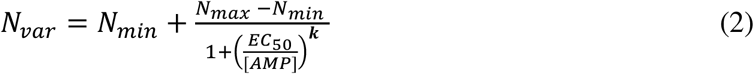

Where *N*_max_ and *N*_min_ correspond to the maximum and minimum value of the variant population size, *N*_var_, [AMP] refers to the AMP concentration in μg/mL and *k* represents the Hill coefficient, which determines the rate of change of the curve.

### Quality of fitness scores and EC_50_ values

Next, we confirmed that our experiments yielded reliable fitness and *EC*_50_ values for the ~5600 variants across a range of AMP conditions. The fitness scores show a strong linear correlation between replicates, with an R^2^ of at least 0.92 for all conditions except at the highest AMP concentration (256 μg/mL), which displayed an R^2^ of 0.45 (**Fig. 1D** and **S2**). The observed noise arising at 256 μg/mL between replicates likely results from the use of an AMP concentration much higher than the *EC*_50-wt_ (84 μg/mL). At this high concentration, the frequency of the *wt*VIM-2 after selection, which is the denominator in **equation (1)**, is drastically reduced and introduces greater error in this condition. The fitness scores show distinct separation between functional and non-functional variants. Variants with synonymous mutations are centered at a fitness score of 0 and the variants containing a stop codon mutation are centered near a fitness of −4 (**Fig. 1D** and **S2**). Hence, a fitness score of −4 or lower denotes inactive variants, and scores that are below −4 are set to −4 in subsequent analyses.

We also found the *EC*_50_ values to strongly correlate between replicates, with a linear regression R^2^ of 0.98 (**Fig. 1E**). Moreover, the *EC*_50_ values derived from DMS are comparable with those measured on individual variants by a conventional dose-response curve resistance assay using the same selection method (87 unique variants, **Data S3**), with a strong linear regression R^2^ of 0.88 (**Fig. 1F**).

### Dependence of fitness on phenotype and selection pressure

The fitness scores we obtained across a range of AMP concentrations allows us to examine how the selection pressure influences the distribution of fitness effects (DFE). The fitness scores reflect relative growth of *E. coli* expressing a given VIM variant relative to *wt*VIM-2, and by extension, the resistance conferred to *E. coli* by a VIM variant in each AMP concentration. The resistance conferred by each variant is further dependent the *EC*_50_, which reflects the variant’s enzymatic efficiency (*k*_cat_/*K*_M_) of and/or the amount of enzyme expressed in the periplasm fraction of the cell. Thus, the DFE is inherently affected by both protein phenotype as well as the external selection environment.

Overall examination shows the DFEs and the distribution of *EC*_50_ values are bimodal (**Fig. 2**), as typically observed in other proteins (*16*, *17*, *20*, *23*, *26*, *27*, *31*–*33*, *44*, *45*). In each distribution, one peak is formed by variants of near-neutral effects and a second peak is formed from variants with strongly deleterious effects, while only a small fraction of mutations exhibit higher values than *wt*. To further quantify and classify the variants in the distributions as having ‘negative’, ‘neutral’ or ‘positive’ effects with respect to the *wt*, we performed two-tailed z-tests between the fitness (or *EC*_50_) of the non-synonymous variants against the distribution of the synonymous variants (a null model with N=251); test results are controlled for false discovery rate by the Benjamini-Hochberg procedure with α = 0.05. The distribution of *EC*_50_ exhibits the most distinct separation between neutral and non-functional variants, where 63% are negative (*EC*_50_ < 72 μg/mL), 35% of mutations are neutral (72 μ g/mL ≤ *EC*_50_ ≥ 171 μg/mL) and 2% are positive (*EC*_50_ >171 μg/mL) (**Fig. 2A**).

**Fig. 2.**
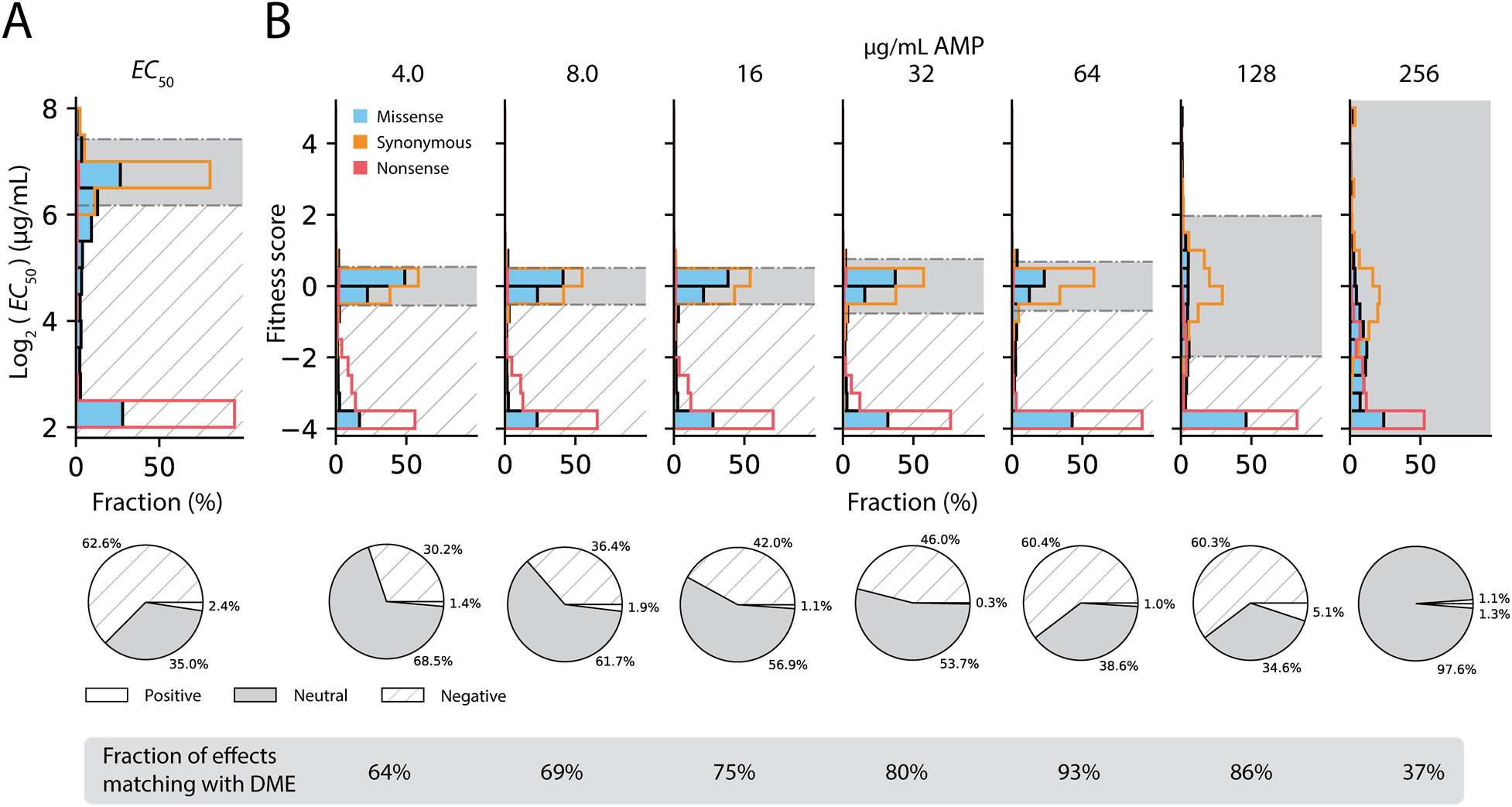
Distributions of fitness and *EC*_50_ effects. (**A**) The distribution of *EC*_50_ for all variants. (**B**) The DFEs of all variants under selection across all AMP selection concentrations. In all plots, the distributions are shown as a percentage of all variants within a mutation type (synonymous, missense or nonsense), with the color legend in **B**). The background colors indicate regions of positive (white), neutral (gray) and negative (hatched) effects, with the legend and a pie chart summary of the proportion of positive/neutral/negative effects below each histogram. For DFEs, the proportion of variants where positive/neutral/negative effects match the effects in the *EC*_50_ are displayed as percentages at the bottom.

Naturally, fitness in the DFEs decline with increasing AMP concentration (**Fig. 2B**), as can be inferred from the overall growth of the *E. coli* transformed with the library (**Fig. 1B**). At lower AMP concentrations of 4.0-32 μg/mL, the majority (54-69%) of variants exhibit neutral effects. When the AMP concentration approaches two-fold of the *EC*_50-wt_ of 84 μg/mL, the majority of variants become negative, up to 60%. At concentrations greater than *EC*_50-wt_ (128-256 μg/mL), the distribution of synonymous mutations becomes broader as the *wt* resistance is also no longer sufficient to sustain high growth, and the data becomes noisier. In particular, at 256 μg/mL almost all variants are neutral again relative to *wt*VIM-2.

The shifting of the DFEs illuminates how mutational responses can be substantially altered depending on the selection pressure. For example, when comparing variant effects (negative/neutral/positive) across the DFEs and the distribution of *EC*_50_ values, only 52.3% of mutations exhibit the same effect across all distributions, with 26.7% consistently deleterious, 25.6% consistently neutral and none that are consistently positive. The remaining 47.7% of mutations exhibit distinct effects dependent on the selection pressure, where 39.5% switch between neutral and deleterious, 5.9% switch between positive and neutral, and 2.1% switch between all three states. We find that effects in the DFEs at 64-128 μg/mL mostly closely resemble the effects in *EC*_50_, where 93% and 86% of variants have the same effect across fitness and *EC*_50_, respectively (**Fig. 2**); as these DFEs are measured at the closest concentrations to the *EC*_50-wt_ of 84 μg/mL, the full extent of negative mutational effects is captured by the fitness scores without affecting the *wt* fitness.

### Phenotype-fitness relationships across different AMP concentrations

Next, we seek to quantify the relationship between the *EC*_50_ values and the fitness scores obtained at each antibiotic concentration. We fit their relationship with a ‘threshold’ model (*9*, *10*, *19*, *20*, *22*, *35*), *i.e.*, the four-parameter sigmoidal function in **equation (3)** (**Fig. 3**).

**Fig. 3.**
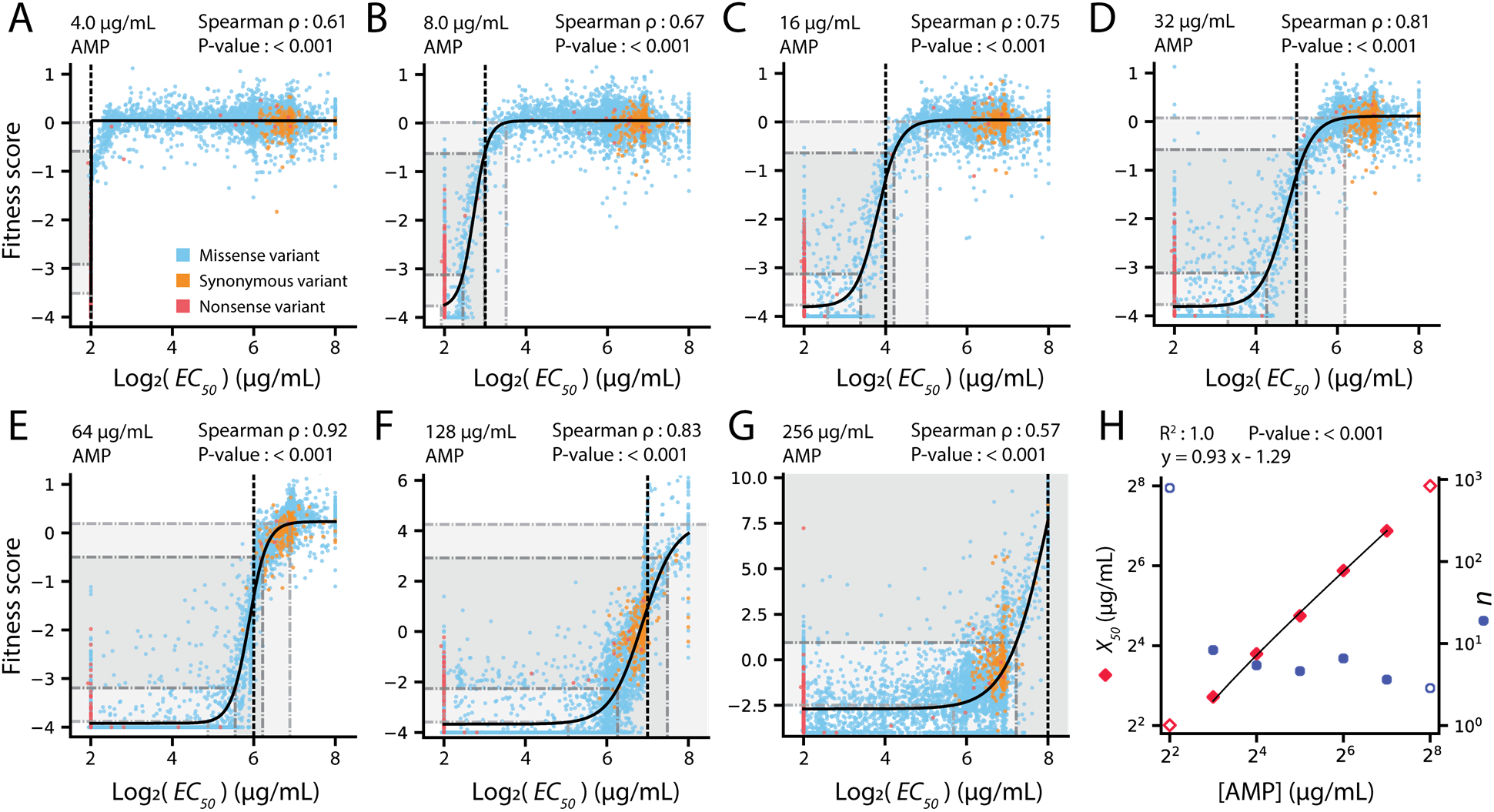
Phenotype-fitness landscapes across different AMP concentrations. Panels **A**-**G** show the comparison of DMS fitness scores with *EC*_50_ of all variants under selection in (**A**) 4.0 (N=5517), (**B**) 8.0 (N=5516), (**C**) 16 (N=5503), (**D**) 32 (N=5517), (**E**) 64 (N=5517), (**F**) 128 (N=5517) and (**G**) 256 (N=5517) μg/mL AMP. In each panel, variants are colored according to mutation type with the color legend in **A**). The solid black curve indicates the line of best fit for a sigmoidal curve fit (**equation 3**), while the black vertical dashed line indicates the AMP concentration used during selection. The dark grey regions extending to the x and y-axes indicates the region of linear association in the sigmoidal curve. The light grey regions extending to both axes indicate the range between 1% - 99% fitness in the fitted curve. The correlation coefficient and P-value of a Spearman rank-order correlation between the fitness score and *EC*_50_ is shown above each plot. (**H**) Sigmoidal fit parameters (**equation 3**) for the curves shown in panels A-G. The *X*_50_ values are shown in orange diamonds and the Hill coefficients (*n*) are shown in blue circles. The filled symbols indicate the conditions where the region of linear association in the sigmoidal curve (dark grey) was fully captured by the AMP concentrations of the experiment, while open symbols indicate the opposite. The R^2^, P-value and equation for the linear regression of *X*_50_ with [AMP] is shown above the plot. Plots of all parameters can be found in **Fig. S3**.

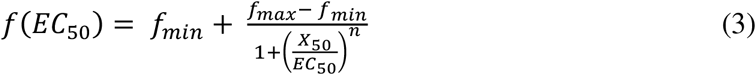

The *f*_min_ and *f*_max_ denote the minimum and maximum fitness scores, respectively, while *n* represents the Hill coefficient. The inflection point of the curve, also the concentration of AMP where fitness is half-way between *f*_min_ and *f*_max_, is denoted by *X*_50_.

Overall, the threshold model recapitulates the phenotype-fitness relationship across all AMP concentrations, with Spearman rank-order correlation between fitness and *EC*_50_ ranging from 0.57-0.92 (**Fig. 3A-3G**). Outside the sigmoidal transition range, fitness scores plateau at high and low fitness for variants with higher and lower *EC*_50_, respectively. In particular, at AMP concentrations of 4.0-64 μg/mL (**Fig. 3A-3E**), *f*_min_ and *f*_max_ maintain steady values in the fit at an average of −3.8 and 0.1, respectively (**Table 1**, **Fig. S3A** and **S3B**). Since, *wt*VIM-2 can fully survive in these conditions, higher *EC*_50_ variants do not exhibit any fitness advantage over *wt*VIM-2. The existence of a fitness plateau for variants with an *EC*_50_ higher than the selection threshold also suggests that there are no systematic trade-offs between *EC*_50_ and fitness in our experimental system; higher *EC*_50_ variants do not exhibit low fitness in the presence of low AMP. At concentrations above *EC*_50-wt_ (128 and 256 μg/mL) the *f*_max_ increases as *wt*VIM-2 becomes challenged and exhibits decreased growth, while higher *EC*_50_ variants continue to grow (**Fig. 3F and 3G**).

**Table 1.** Fitted parameters of sigmoidal phenotype-fitness relationships.

| [AMP]<br>(μg/mL) | Equation fit <sup>a</sup> |  |  |  | Linear range <sup>a</sup> |  |  |  | Transition range <sup>a</sup> |  |  |  |
| --- | --- | --- | --- | --- | --- | --- | --- | --- | --- | --- | --- | --- |
| | $f_{\max}$ | $f_{\min}$ | $n$ | $X_{50}$<br>(μg/mL) | $EC_{50}$ lower<br>(μg/mL) | $EC_{50}$ upper<br>(μg/mL) | $f_{\text{lower}}$ | $f_{\text{upper}}$ | $X_1$<br>(μg/mL) | $X_{99}$<br>(μg/mL) | $f_{1\%}$ | $f_{99\%}$ |
| 4 | 0.0 | -3.5 | 788 | 4.0 | 4.0 | 4.0 | -2.9 | -0.6 | 4.0 | 4.0 | -3.5 | 0.0 |
| 8 | 0.0 | -3.8 | 8.3 | 6.6 | 5.5 | 7.9 | -3.1 | -0.6 | 3.8 | 11.4 | -3.8 | 0.0 |
| 16 | 0.0 | -3.8 | 5.4 | 13.9 | 10.5 | 18.5 | -3.1 | -0.6 | 6.0 | 32.4 | -3.8 | 0.0 |
| 32 | 0.1 | -3.8 | 4.6 | 26.9 | 19.2 | 37.5 | -3.1 | -0.6 | 9.9 | 72.7 | -3.8 | 0.1 |
| 64 | 0.2 | -3.9 | 6.6 | 58.9 | 46.6 | 74.5 | -3.2 | -0.5 | 29.3 | 118 | -3.9 | 0 |
| 128 | 4.3 | -3.7 | 3.6 | 117 | 76.7 | 179 | -2.3 | 2.9 | 33.2 | 415 | -3.6 | 4 |
| 256 | 18.0 | -2.7 | 2.9 | 256 | 149 | 439 | 0.9 | 14.4 | 51.4 | 1280 | -2.5 | 18 |

<sup>a</sup> All parameters shown are calculated from a fit to equation (3).

The quantification of the phenotype-fitness relationships reveals some notable traits. First, the transitions of fitness between the lower and upper fitness plateaus (*X*_1_- *X*_99_, points in the curve at 1% and 99% fitness) (**Table 1**) occur in a relatively small range of *EC*_50_ differences (3.0-7.3 fold range) when considering conditions where the transition can be fully resolved within experimental conditions (8.0-64 μg/mL AMP). This confirms AMP selects the phenotype of VIM-2 variants in a discrete, threshold-like manner. Second, where the linear range can be fully captured (8.0-128 μg/mL AMP), the shape of the transitions occur in a very similar fashion across AMP concentrations; the Hill coefficient remaining steady with an average of 5.7 ± 1.8 S.D, leading to linear transition ranges that spans a 1.4 - 2.3 fold change in *EC*_50_. (**Fig. 3H, S3C** and **S3D**). This suggests that the mode of selection by AMP against VIM phenotypes is universal across a wide range of AMP concentrations. Third, the inflection points (*X*_50_) in the phenotype-fitness relationship are a nearly perfect function of AMP concentration as described by the linear relationship in **equation (4)** (**Fig. 3H**).

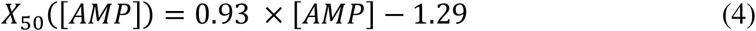

Taken together, the selection of VIM-2 phenotypes is strongly dictated by the concentration of AMP in the environment, and the range of phenotypic selection is largely predictable across a defined range of concentrations.

### A phenotype-environment-fitness landscape of VIM-2

Utilizing the universal behavior of the Hill coefficient (*n*) and the inflection points (*X*_50_) in the phenotype-fitness relationships, we propose a simple, empirical model to describe the complete phenotype-fitness landscape of VIM-2 under AMP selection in **equation (5)**.

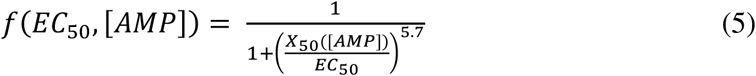

This equation describes the fitness (*f*) of a variant as a function of its *EC*_50_ and the AMP concentration. All parameters are functionally the same as **equation (3)**, but *n* = 5.7, and the inflection point of the curve (*X*_50_) is now a linear function of [AMP] as shown in **equation (4)**. Furthermore, we normalize the observed fitness scores to the *f*_min_ and *f*_max_ determined from the sigmoidal fits at each AMP concentration in **Table 1**, such that a fitness of 0 represents the minimum fitness and a fitness of 1 represents the maximum fitness (**Fig. 4A**). The model enables us to depict the entire phenotype-environment-fitness landscape, which shows agreement with ~39,000 data points across seven different selection conditions (**Fig. 4B**). Thus, the universal description by a simple mathematical model suggests that the selection of VIM variants is highly dictated by the environmental AMP concentration, and that the evolutionary response of any given variant can be rather predictable.

**Fig. 4.**
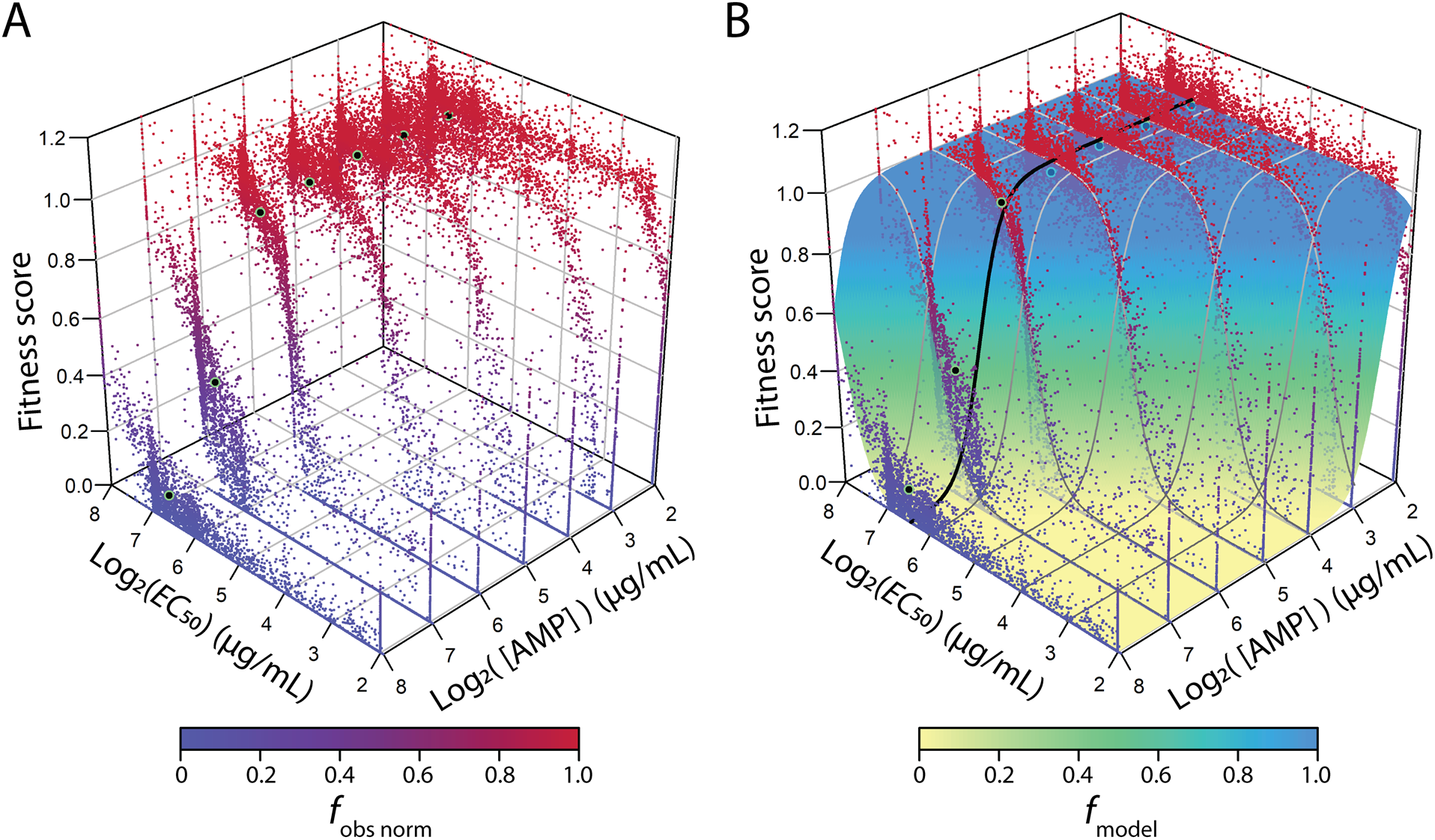
The VIM-2 Phenotype-fitness landscape as a function of *EC*_50_ and AMP concentration. (**A**) For each variant, the observed fitness score is plotted in relation to the AMP concentration during selection, and its respective *EC*_50_. The fitness scores are normalized at each concentration according to **equation (13)** in the Methods. The black dots represent the fitness of *wt*VIM-2 after normalization. (**B**) The modelled behavior of fitness scores according to **equation (5)**. The dots are the same data as panel **A**) while the surface depicts the predicted fitness scores according to the model. The black line indicates the model fitness at the *EC*_50-wt_. Cross sections of the model at two-fold intervals of *EC*_50_ or AMP concentrations are marked by grey lines.

### Insight into the evolutionary dynamics of sub-MIC selection

Our global phenotype-environment-fitness landscape allows us to gain quantitative insights into an important question regarding the evolution of antibiotic resistance: a phenomenon known as sub-MIC selection. Antibiotic concentrations well below the ‘minimal inhibitory concentration’ (MIC) of the *wt* in the population can still promote the emergence of higher resistance variants (*46*–*50*). Typically, any resistant variant with *MIC*_var_ higher than *MIC*_wt_ will have an absolute resistance and fitness advantage when the antibiotic concentration is over *MICwt* (*i.e.*, when *MIC*_wt_ < [AMP] < *MIC*_var_) (*46*, *51*–*54*) (**Fig. 5A**). In sub-MIC selection where the antibiotic concentration is below *MICwt*, the *wt* is still viable, but the resistance difference could still lead to the selection of the variant over the *wt*, provided the benefit of resistance is high enough to offset any trade-offs due to fitness costs (*46*, *51*). The lowest antibiotic concentration where the variant can overtake the *wt* in fitness is designated the ‘minimal selective concentration’ or MSC. Quantitative knowledge supporting existing models of sub-MIC selection remains rare, however (*46*, *55*).

Our model can bridge this gap by providing a quantitative depiction of the mechanism underlying sub-MIC selection. For example, *wt*VIM-2, with *EC*_50_ of 84 μg/mL, would exhibit an MIC (1% of *f*_max_) of 203 μg/mL AMP, and an MSC (99% of *f*_max_) of 42 μg/mL (see ‘**Calculation of MIC and MSC**’ in methods). Hence, for *wt*VIM-2, the range for sub-MIC selection can extend to ~5-fold below the MIC*wt*. Furthermore, we can calculate the fitness difference between *wt*VIM-2 and variants exhibiting higher resistance (**Fig. 5B**). We observe that selection can still strongly favor the higher resistance variant (*f*_var_ - *f*_wt_ > 0.1) until the AMP concentration drops to ~2-fold above the MSC (**Fig. 5B**). This is the case even when the variant is only 1.5 to 2-fold more resistant than the *wt*, with the fitness difference being more pronounced for variants with higher resistance. Thus, AMP concentrations as low as 5-fold below the *wt* MIC can still select for variants with higher resistance than *wt*VIM-2, which is largely consistent with studies of sub-MIC selection using other distinct systems (*46*–*50*).

**Fig. 5.**
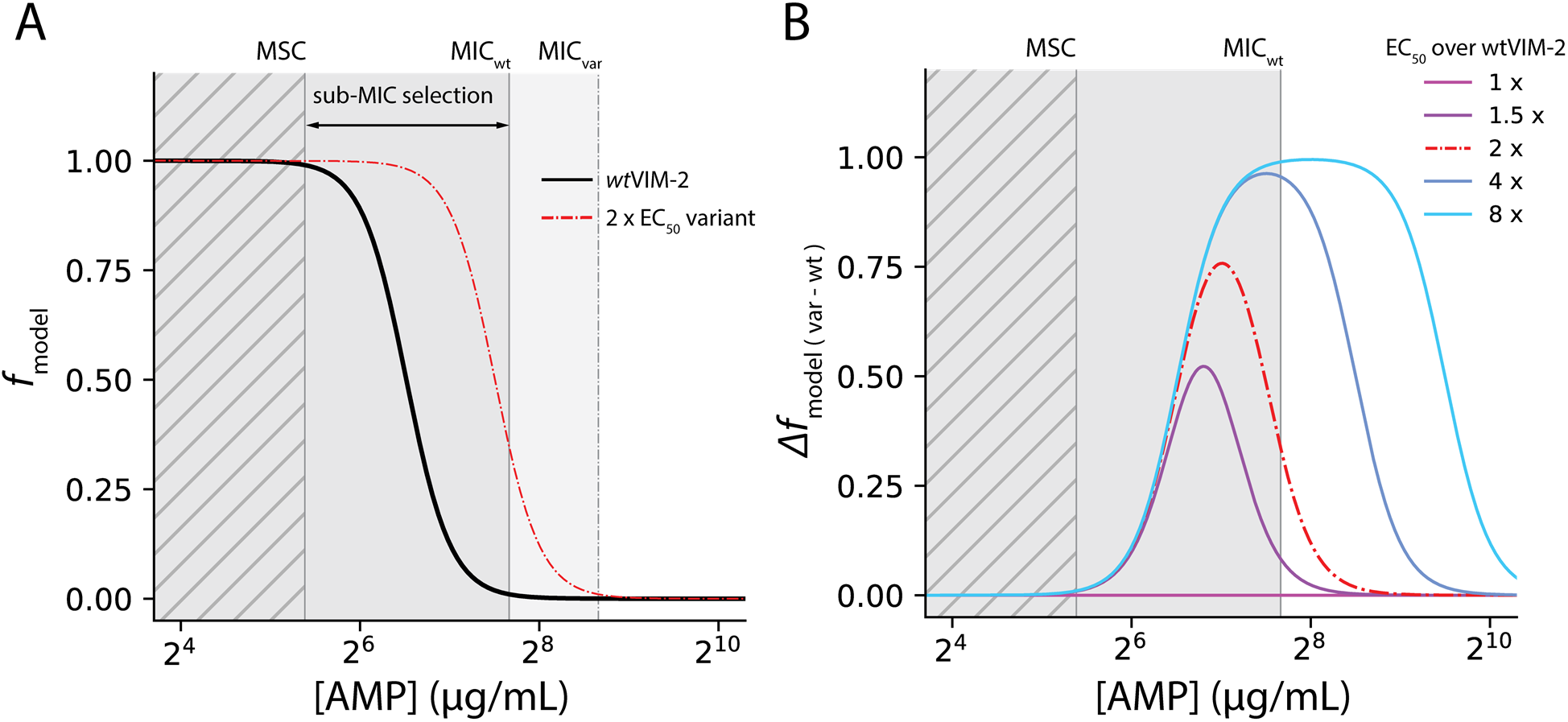
Sub-MIC selection behavior of VIM-2 variants. (**A**) The fitness of *wt*VIM-2 and a hypothetical variant with 2 × *EC*_50_-_wt_ as a function of AMP concentration at a fixed *EC*_50_. The MIC (*f* = 0.01) of both curves and the MSC (*f* wt = 0.99) are marked as vertical lines. The region in light grey is the ‘mutant selection window’, the region in dark grey is the ‘sub-MIC selection window’ and the hatched region indicates no selection of the variant over wt. (**B**) The difference in fitness between *wt*VIM-2 and higher resistance variants across different AMP concentrations. Each curve indicates a variant with an *EC*_50_ of 1-8 fold relative to *wt*VIM-2. The coloring of the sub-MIC selection range in the background is the same as in **A**).

### The molecular basis for nonspecific epistasis

Our empirical model can also provide a quantitative understanding of the mechanisms underlying mutational epistasis, *i.e*., the non-additive interaction between mutational effects (*56*, *57*). Epistasis is a key phenomenon in protein evolution that dictates the shape and ruggedness of fitness landscapes, which subsequently affect important evolutionary phenomenon such as robustness (*9*, *11*, *12*, *15*, *18*, *19*), evolvability (*6*, *10*, *12*, *13*, *20*–*22*, *58*) and evolutionary contingency (*29*, *56*, *59*). One form of epistasis, defined as ‘nonspecific epistasis’, is known to be highly prevalent and strongly influences the accessibility of mutations. Nonspecific epistasis typically stems from the non-linear relationship between fitness and protein phenotype, such as we observed between fitness and *EC*_50_ in this study (**Fig. 3**). The key distinction of nonspecific (as opposed to specific) epistasis is that epistasis can still be observed at the level of fitness even if the mutational effects of two mutations on a phenotype, *i.e.*, *EC*_50_ (and thus enzymatic efficiency or protein expression) are non-epistatic (changes relative to *wt* are log-additive, or more simply described as ‘additive’) (*6*, *14*, *15*, *17*, *18*, *56*, *57*, *60*). While the existence of nonspecific epistasis is widely appreciated, we lack quantitative understanding of the extents to which different types of epistasis can arise depending on the selective environment.

To this end, we simulated the prevalence and types of nonspecific epistasis in VIM-2 fitness using the empirically measured distribution of *EC*_50_ and the phenotype-environment-fitness landscape (**Fig. 6**). Since each variant only contains a single mutation, we can treat the variants’ effects directly as mutational effects. Specifically, we randomly sampled 200 single point missense mutations, calculated the Δ*EC*_50_ values of a given mutation from *wt* (**Fig. 6B**), then combined all pairwise mutations based on a null model (**Fig. 6C**)—a double mutational effect on *EC*_50_ is equal to the additive effect of two single mutations. All pairwise combinations of 200 mutations leads to _200_C_2_ = 19900 mutant pairs. Then, using the model in **equation (5)**, we computed the change in fitness (Δ*f*) for all individual and pairwise combinations at each AMP concentration using the single and pairwise *EC*_50_. We quantified epistasis by comparing the observed fitness change (Δ*f_obs_*) calculated from the pairwise *EC*_50_, and the expected fitness change (Δ*f_exp_*), which is the sum of the Δ*f* of both individual mutations (**Fig. 6D**, See methods for ‘**Nonspecific epistasis of VIM-2 variants**’). Finally, we categorized the observed epistasis into ‘positive’ or ‘negative’, based on the observed fitness change compared to the expected additive change (*56*) (**Fig. 6A**). Positive epistasis is where Δ*f_obs_* is more positive than Δ*f_exp_* and negative epstasis is where Δ*f_obs_* is more negative than Δ*f_exp_*. We also take note of cases where Δ*f_obs_* has an opposite sign to Δ*f_exp_*.

**Fig. 6.**
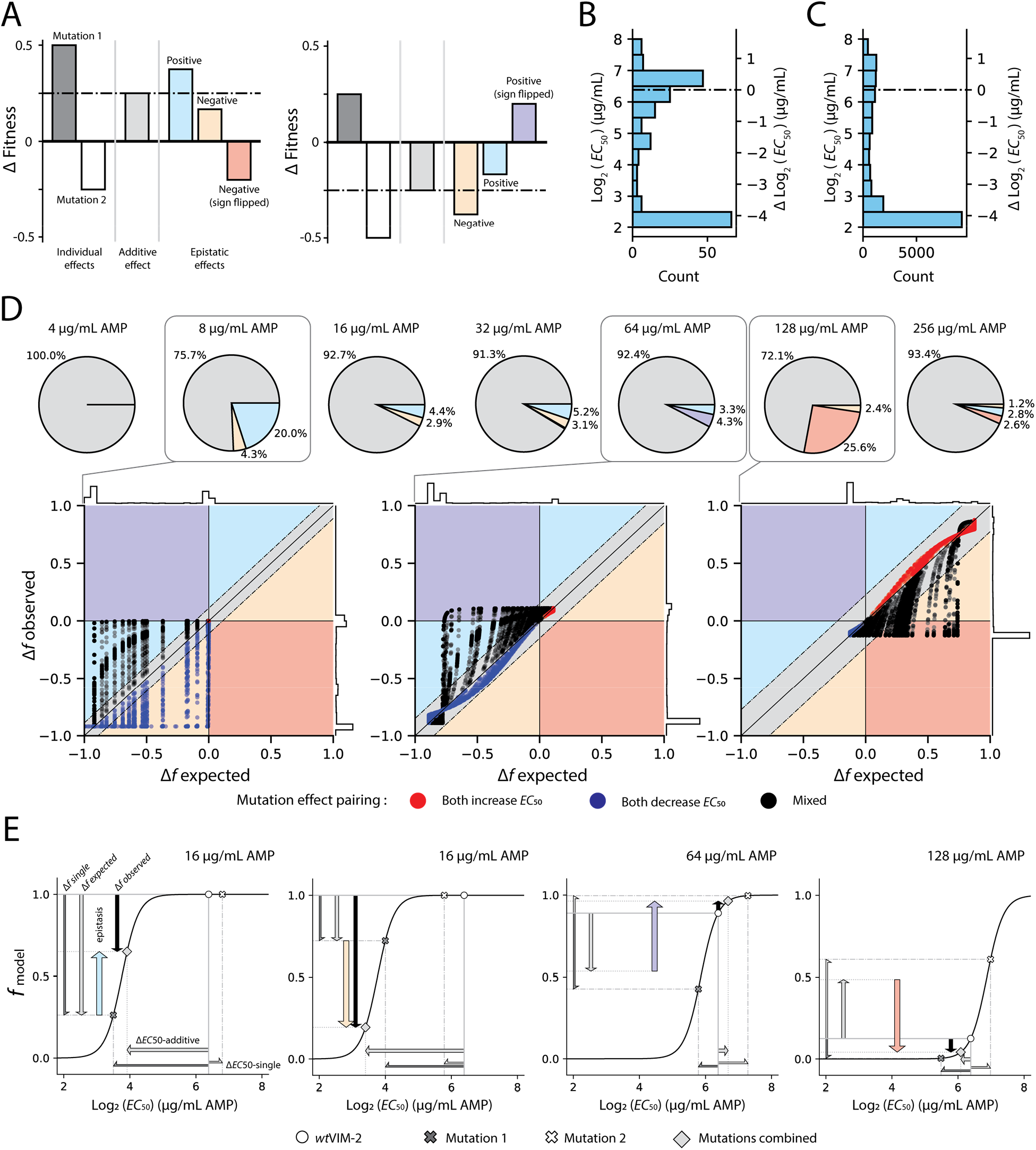
Nonspecific epistasis varies across selection conditions. (**A**) Conceptual illustration of each type of epistasis considering an overall positive (left) or negative (right) additive effect. This given color scheme for epistasis types is maintained throughout the figure. (**B**) Distribution of the 200 randomly sampled *EC*_50_ values, with the change in *EC*_50_ relative to *wt*VIM-2 displayed on the right y-axis. (**C**) Distribution of the 19900 pairwise additive *EC*_50_ values, with the change in *EC*_50_ relative to *wt*VIM-2 displayed on the right y-axis. (**D**) The proportion of mutational pairs exhibiting epistatic effects under each condition; only proportions higher than 0.5% are labelled. Below the pie charts are scatterplots from key conditions showing the difference between expected and observed fitness changes. The backgrounds of the scatterplots are also colored according to the type of epistatic effect. A gray background indicates no significant epistasis, or additivity, and is defined as a range of 1.96 × S.D. of the synonymous mutant fitness distribution after normalization by **Equation (13)** for each AMP concentration. In each scatterplot, points representing the combined effect of a pair of mutations are colored according to the effect of each single mutation (red when both increase *EC*_50_ over *wt*VIM-2, blue when both decrease *EC*_50_ and black when one increases while the other decreases *EC*_50_). Histograms of all data points in each scatterplot are shown for each axis. (**E**) Visual depictions of each type of epistasis at distinct AMP concentrations, as a result of a non-linear relationship between fitness and *EC*_50_.

The proportion of each type of epistasis is drastically dependent on the environment (0 - 28% epistasis). At the lowest AMP concentration (4 μg/mL), no epistasis can be observed as most mutations simply exhibit neutral effects. At 8 μg/mL AMP, the greatest proportion of positive epistasis is observed (20%), and about 4% of combinations showing negative epistasis. At 16 and 32 μg/mL AMP, smaller fractions of positive and negative epistasis are observed (7.3% and 8.3% in total, respectively). From 64-256 μg/mL AMP, a substantial fraction of mutational combinations exhibit epistasis strong enough to flip the sign of the double mutant fitness effect. A small fraction (4.3%) of expected negative fitness effects are ‘flipped’ into positive observed effects at 64 μg/mL AMP, while a striking 26% of expected positive fitness effects are flipped into negative observed effects at 128 μ g/mL AMP. It should be emphasized that these substantially different epistasis patterns observed in different environments are based on the same set of mutational combinations within the experimentally obtained distribution of *EC*_50_, even with non-epistatic effects on the phenotypic level (*EC*_50_); such patterns are the results of a non-linear mapping between phenotype and fitness.

An examination of the non-linear relationship can provide the mechanistic explanation underlying the observed nonspecific epistasis, and note a certain degree of predictability in the type of epistasis observed (**Fig. 6E**). Key to the phenomenon is the location of the final combined phenotype relative to the sigmoidal fitness transition (a threshold that is determined by the selection pressure). Positive epistasis is highly prevalent at lower selection pressures (8 μg/mL AMP) as even small increases in phenotype can be enough to ‘climb’ the nearby fitness transition, rescuing the fitness of seemingly non-functional variants. Similarly, substantial negative epistasis occurs at selection pressures just above the *EC*_50-wt_ of 84 μg/mL (128 μg/mL AMP) as small decreases in phenotype can cause neutral and mildly positive variants to rapidly ‘fall off’ the fitness transition. The proportion of epistasis is dependent on the density of variants that are near the selection threshold, and are thus more likely to traverse the fitness transition, in the underlying *EC*_50_ distribution (**Fig. 2A**). Sign flipping is only present near AMP concentrations close to the *EC*_50-wt_ of 84 μg/mL, and occurs only between mutant combinations of opposite effects; sign flipping cannot occur when the *wt*VIM-2 is on a high or low fitness plateau, where fitness changes can only occur in one direction (*e.g.*, fitness can only decrease when on the high fitness plateau). Positive sign flipping is only observed when [AMP] is below *EC*_50-wt_ while negative sign flipping only occurs when [AMP] is above *EC*_50-wt_ (**Fig. 6D**).

Taken together, using the empirical distribution of mutational effects and the threshold model that we have established, we demonstrated that the selective environment strongly affects the resulting epistasis. We have also illustrated how epistasis can be prevalent in largescale mutational studies using fitness measurement (*13*, *16*, *17*, *23*, *60*–*63*), as epistasis can still arise even when the underlying phenotype simply experience an additive change (*14*). Notably, we demonstrate a certain degree of predictability in the patterns of nonspecific epistasis: if two mutations have a combined additive phenotype that approaches the selection threshold, one can expect deviations from additivity in fitness. As the trend of apparent epistasis can be drastically different according to the environment, an understanding of the environmental factors is essential to decipher the evolutionary dynamics of proteins. Note that, in this model, we assumed additive effects on *EC*_50_, while empirical mutational combinations are likely to exhibit non-additive effects with respect to protein function and stability due to specific epistasis (*56*, *57*, *64*). In such cases, epistasis will be even more frequently observed, and a rigorous model such as ours may help to distinguish between nonspecific and specific epistasis within high-throughput mutational experiments.

## Discussion

In this study, we have presented a phenotype-environment-fitness landscape of VIM-2 β-lactamase, constructed using a comprehensive set of ~39,000 data points, across a 64-fold range of AMP concentrations. Our observations provide a long-sought, quantitative verification of an important model for protein evolution, the ‘threshold’ model. With this comprehensive and empirical dataset, we unveil how the phenotype-fitness relationship can be shaped by environmental variation, particularly the selection strength. Importantly, we demonstrate that a simple quantitative model can recapitulate the behavior of VIM-2 β-lactamase within the landscape. Our findings provide evidence that the evolution of resistance in a population follows a relatively simple and predictable rule, dictated through the selection pressure exerted by β-lactam antibiotics. Moreover, we provide a mechanistic understanding of how different environments can cause different types of epistasis to arise, solely from knowledge of the threshold relationship between protein phenotype and fitness. Our observations suggest that the influence of the environment on the phenotype-fitness relationship is extremely important to understand the evolutionary dynamics of proteins (*55*, *65*). Hence, strong understanding of the effect of environmental factors may allow us to accurately predict the evolution of antibiotic resistance genes and other proteins.

Since we employed VIM-2, a class B β-lactamase, as a model for this study and measured changes in *EC*_50_—an important trait commonly measured trait for β-lactamases as well as other antibiotic resistance genes—our model may be applicable to other β-lactamases, even in other classes (A, C and D classes). Our findings could prove useful in developing a more general understanding of β-lactamase evolution. Indeed, our observations agree with several behaviors observed in β-lactamases, or other antibiotic resistance genes, beyond just the threshold-like selection behavior. The ~5-fold range of sub-MIC selection estimated from our model largely matches with other studies of the sub-MIC selection range in other systems (*46*–*50*). We do not observe any notable, systematic trends that indicate trade-offs between high *EC*_50_ and fitness at lower AMP concentrations, supporting observations that higher resistance does not necessarily convey a large fitness cost, and that it can be maintained even under low selection pressure (*46*, *47*). This also matches previous modelling where a flat maximum fitness plateau is found for the mutants of TEM-1 irrespective of the selection pressure (*20*). The lack of a fitness cost for more resistant variants is likely a result of the non-essential nature of VIM-2 in our model system, coupled with a fairly moderate protein expression level. Of course, when present, fitness trade-offs add greater complexity to the dynamics of sub-MIC selection (*46*, *55*). Regardless, our quantitative descriptions for those phenomena are invaluable to further deepening our understanding of the behaviors of these antibiotic resistance genes.

We note our relatively simple experimental system poses limits on the generalization of our findings, and additional complexities may be studied in the future. *i*) We only mutated within the VIM-2 gene, and mutations in other genetic context such as the promoter and origin of replication of the plasmid might trigger different dynamics. *ii*) We conducted selection over 6 hours of continuous growth, up to the point where the cells are reaching the early phases of growth saturation. Different culturing conditions may also expose different dynamics. *iii*) The results of AMP selection in *E. coli* may not reflect the behavior of VIM-2 with other classes of antibiotics, such as cephalosporins or carbapenems, as well as other clinically important bacterial hosts, *e.g.*, *P. aeruginosa* and *A. baumannii*. *iv*) These findings from a simple laboratory environment may only be partly applicable to behavior in natural environments, which will have additional variation outside of antibiotic concentration. *v*) Though we observed no systematic fitness-resistance trade-offs, higher resolution measurements of individual variants, and comparison to a condition without any selection may reveal trade-offs that are not as pronounced in our system (*21*, *46*, *47*, *49*, *66*).

Nonetheless, we demonstrated that a quantitative mapping of the fitness landscape using deep sequencing is an extremely powerful method to study protein evolution. Similar studies for more complex conditions, and of different model proteins, will be key for further developing our understanding of protein evolution, in particular, the molecular mechanisms underlying key evolutionary phenomenon like mutational robustness, epistasis and evolvability. Ultimately, better knowledge of phenotype-fitness relationships will also improve our ability to predict the evolution of proteins, such as antibiotic resistance genes, in a complex, natural environment. Such knowledge should also lead to better design and engineering strategies to generate novel proteins and metabolic pathways.

## Acknowledgements

We thank Adrian Serohijos, Pouria Dasmeh and members of the Tokuriki lab for insightful comments on the manuscript.

## Author contributions

John Z Chen: Conceptualization, Software, Formal analysis, Investigation, Methodology, Writing - original draft, Writing - review and editing; Douglas M Fowler: Software, Validation, Methodology, Writing - review and editing; Nobuhiko Tokuriki: Conceptualization, Supervision, Funding acquisition, Project administration, Writing - review and editing

## Data and materials availability

All data is available in the manuscript or the supplementary materials.

## List of Supplementary Materials

Materials and Methods Figs. S1-S3

References (*67*, *68*)

Data S1-S3

## Materials and Methods

### Deep mutational scanning of VIM-2

The variant library of VIM-2 was generated in a previous study, and all selection experiments, deep sequencing and quality control for sequencing data were conducted in the same manner as before(*33*). Briefly, the plasmid based wild-type VIM-2 gene (*wt*VIM-2) gene was mutated into a library containing all single amino acid substitutions through PCR mutagenesis using a primer that replaces each codon position with a degenerate ‘NNN’ sequence. The library of all 267 codon positions was divided into 7 groups of 39 amino acids (31 in the last group), forming 117nt ‘tiles’ that can be sequenced with full overlapping reads using Illumina Next-seq 550. The libraries were transformed into *E. coli* E. cloni 10G electrocompetent cells (Lucigen) using the accompanying protocol, and grown in liquid culture (Miller LB broth) without selection, or selected with 4.0-256 μg/mL AMP in two-fold increments for 6 hours at 37°C. After 6 hours the cell concentration of the *E. coli* culture was measured by optical density at 600 nm (*OD*_600_). The culture was then removed from selection by centrifugation and aspiration of the selection media, then resuspended in fresh LB; this procedure was repeated for a total of three washes. The washed cells were grown overnight, then the plasmid DNA was isolated using a QiaPrep 96 DNA extraction kit (Qiagen). One ‘sample’ in this experiment encompasses a single replicate of one library group of variants grown under a single condition. This process was conducted in two replicates on separate days, while an additional replicate (three total) was performed for the non-selected library. The third non-selected library was initially intended as a control to detect if variants in the library are lost during the 6 hours of growth without selection, and was grown overnight but was not subject to the 6 hours of growth at 37°C. However, we observed no notable differences in variants present between the three replicates of the non-selected library and the third replicate was treated the same as the other two replicates in subsequent analysis.

Following selection and DNA extraction, plasmid DNA from each library group was amplified by PCR for 15 cycles, using primers that amplify only the 117 bp region of each group and attach the universal Nextera adaptor sequence to each end of the amplicon. The amplicons are extended by PCR again to include the sample indices (i7 and i5) and flow cell binding sequence, then sequenced using a NextSeq 550 sequencing system (Illumina, Inc.). The *wt*VIM-2 plasmid was also included in the sequencing run to measure deep sequencing error rates. The raw sequencing data can be found in the NCBI Sequencing Read Archive (SRA) (BioProject accession: **PRJNA634597**). Subsequent processing of deep sequencing data and calculation of fitness scores are conducted using our previously published in-house Python scripts (https://github.com/johnchen93/DMS-FastQ-processing). Using these scripts after sequencing, the forward and reverse paired-end sequencing reads are merged and the posterior quality score of each position is calculate from the quality scores of the two reads (*65*). Sequences that have more than 20 mismatches between forward and reverse reads, or contains any position with a posterior quality score less than 10 were discarded. The median number of reads that pass the merging process is ~660,000. Only one of the samples from the non-selected library and 7 samples from the selected conditions had less than 100,000 reads after filtering (all limited to replicate 1), and these low read samples were handled accordingly in the subsequent applications (see sectionsbelow on ‘**Calculation of fitness scores**’ and ‘**Population size estimates using deep sequencing data**’).

The merged sequences were used to identify mutations, and sequences with more than one codon mutated were discarded. To further ensure the quality of the variants we identified, the sequencing errors observed when sequencing *wt*VIM-2 was used to calculate the expected frequency of single amino acid variants arising from sequencing errors alone. Then, the non-selected library was filtered so that variants with frequencies lower than 2 × of the expected frequency from sequencing errors or have a read count less than 5 are excluded from further analysis. Only the variants that pass all of the above quality control measures are used for further calculations. The frequencies of each amino acid variant were calculated using **equation (6)** for each variant. Variants that passed quality filtering in the non-selected library but were no longer present in the selected libraries were given a dummy count of 1 to estimate the lowest possible frequency.

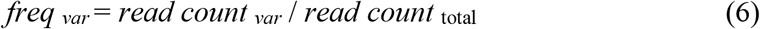

### Calculation of fitness scores

The variant frequencies were used to calculate fitness scores for selection conducted at each AMP concentration using **equation (1)**, where each selected library is matched with the non-selected replicate that was grown in the same replicate for calculations; the extra non-selected library replicate was not considered. Fitness scores were averaged by taking the arithmetic mean and averaged scores below −4.0 were set to a minimum of −4.0 for further analysis. The non-selected library in group 1 of replicate 1 contained less than 10,000 reads after merging and only ~25% of the variants were detected after filtering, meaning that most variants in positions 1-39 only have fitness scores from replicate 2. Additionally, of the 7 samples selected with AMP that contained less than 100,000 reads after FastQ merging, 2 samples were retained after scoring as they share correlation R^2^ greater than 0.8 with the other replicate while the scores from the other 5 samples were excluded; this includes library group 2 (selected with 16 and 128 μg/mL AMP), 4 (8 μg/mL AMP), 6 (16 μg/mL AMP) and 7 (128 μg/mL AMP) from replicate 1. Replicate fitness scores were combined by taking the arithmetic average (**Data S1**).

### Population size estimates using deep sequencing data

Population sizes were calculated using the variant frequencies and the *OD*_600_ of the library after selection. The code used to calculate the population size and *EC*_50_ can be found at https://github.com/johnchen93/DMS-EC50. To consolidate the 3 replicates of the non-selected library, the frequency of each variant was averaged between any replicates where the variant passed quality filtering; frequencies from selected libraries were kept as separate replicates. The frequencies of each variant were then used to estimate the population size of cells containing that variant by multiplying the frequency with the *OD*_600_ observed immediately after the 6 hours of selection, then multiplied again with an estimated cell count of 10^9^ cells per 1 unit of *OD*_600_ using **equation (7)**; all cultures were selected at the same total volume of 1mL.

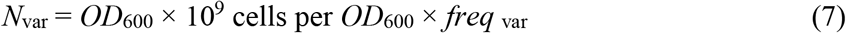

It is possible for the frequency to represent an unrealistic cell count, but in our case even purged variants that have been given a dummy count of 1 will still have population estimates of at least 300-500 cells. Samples with less than 100,000 reads after merging were excluded from this analysis, which consist of library groups 2 (at 16 and 128 μg/mL AMP), 4 (8 μg/mL AMP), 6 (8, 16 and 32 μg/mL AMP) and 7 (128 μg/mL AMP) in replicate 1.

### Calculation of EC_50_ for each variant

The estimated population size of each variant (y-axis) is plotted against the AMP concentration (x-axis) to generate a dose-response curve. A sigmoidal fit is applied to the curve using the scipy.optimize package’s ‘curve_fit’ function to calculate the *EC*_50_ using **equation (2)**. In the initial fit, <u>*N*</u>max is estimated as the average population size of the variant in the non-selected libraries or the library selected at the lowest AMP concentration, whichever is the higher value. *N*_min_, *k* and *EC*_50_ were estimated as 0, −1 and 32μg/mL AMP, respectively. The fitting was performed separately for each replicate.

All dose-response curves were plotted regardless of whether a successful fit was made and each curve fit for each replicate was manually curated. Curves where the *EC*_50_ lies in a region with data missing in two consecutive concentrations are discarded. Some variants exhibit *EC*_50_ transitions outside the measured range of concentrations (4 - 256 μg/mL AMP), in which case the *EC*_50_ is estimated as the lowest (4 μg/mL AMP) and highest AMP concentration (256 μg/mL AMP) if the *EC*_50_ transition occurs past the lowest and highest AMP concentration, respectively. To merge the *EC*_50_ between replicates, we took the arithmetic average of the log2 (*EC*_50_) (**Data S2**).

### Measuring EC50 using conventional drug-response assays

Previously, we measured the *EC*_50_ for AMP of 39 amino acid variants that were isolated and assayed individually(*33*). In this study, we determined the *EC*_50_ of an additional 48 amino acid variants (63 unique codons) and measured their *EC*_50_ using the same method. Variant libraries at positions across the length of the VIM-2 sequence (positions 5, 16, 42, 73, 96, 153, 162, 173, 201, 209 and 251) were transformed into *E. coli*, plated on LB-agar plates and single colonies were picked, cultured and stored in glycerol stocks in a 96 well plate; for each position, we picked 8 colonies. Two colonies each of *E. coli* transformed with *wt*VIM-2 or empty vector were also isolated in the same manner and placed in the same plate as controls. The identity of each isolated variant was confirmed through Sanger sequencing.

Antibiotic selection was conducted using the same procedure as the DMS selection presented above, where a liquid culture of the E. coli containing each isolated variant is grown for 6 hrs under selection with 2.0-256 ug/mL AMP at 2-fold increments, at 37°C. The *OD*_600_ of each culture is measured after selection and plotted against the AMP concentration to generate a dose-response curve. The curve was fitted with the sigmoidal **equation (2)** to calculate the *EC*_50_, where *OD*_600_ was used directly in place of *N*_var_, while *N*_min_ and *N*_max_ were estimated as 0 and the *OD*_600_ from growth without AMP, respectively. Final *EC*_50_ values of the variants picked in this study can be found in **Data S3**.

### Phenotype-fitness relationship at single AMP concentrations

The fitness scores (y-axis) at each AMP concentration were plotted against the *EC*_50_ (x-axis). The relationship between fitness scores and *EC*_50_ at each AMP concentration was fitted to the sigmoidal curve in **equation (3)**. The initial estimates for *f*_max_, *f*_min_ and *n* are 2.0, −4.0 and 1.0 respectively for all AMP concentrations. The initial estimate for *X*_50_, the inflection point of the curve, was set to be the same as the AMP concentration from which the dataset was measured.

The boundaries of the transition between the lower and upper fitness plateaus (*X*_1_-*X*_99_) were calculated as the *EC*_50_ values at 1% and 99% fitness relative to the range between the fitted *f*_max_ and *f*_min_ of each AMP concentration.

The linear region of the sigmoidal curve for the DMS fitness scores was calculated using **equations (8-11)**, based on the final fitted values for each AMP concentration (*66*).

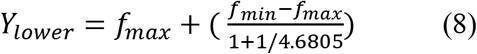

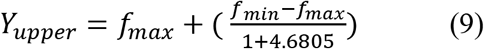

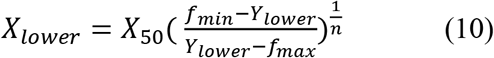

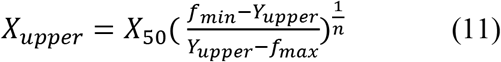

In this case, *X*_lower_ and *X*_upper_ indicate the start and the end of the linear range in terms of *EC*_50_ while *Y*_lower_ and *Y*_upper_ indicate the start and end of the linear range in terms of fitness score.

### General model for the phenotype-fitness landscape of VIM-2

To generate the model, we start by connecting the sigmoidal behavior of the phenotype-fitness relationships across different AMP concentrations into the general form in **equation (12)**.

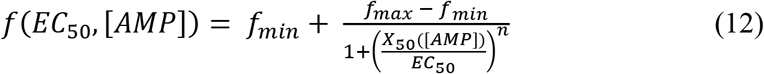

All parameters are functionally the same as **equation (3)** except the inflection point of the curve (*X*_50_), which is now a linear function of AMP as shown in **equation (4)**. The Hill coefficient (*n*) is 5.7, the average where the linear range of the sigmoidal transition was not limited by the selection concentration (8-128 μg/mL). Furthermore, we normalize the observed fitness scores to the *f*_min_ and *f*_max_ determined from the sigmoidal fits at each AMP concentration in **Table 1** using **equation (13)**, such that a fitness of 0 represents the minimum fitness and a fitness of 1 represents the maximum fitness.

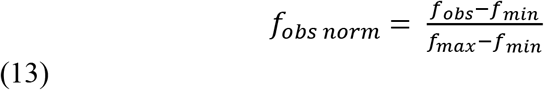

This allows us to arrive at the final form of the model presented in **equation (5)**.

### Calculation of MIC and MSC from the general model

We calculated the minimal inhibitory concentration (MIC) of *wt*VIM-2 as the AMP concentration at which fitness is reduced to a threshold *t* = 0.01 (1% of *f*_max_), according to **equation (14)** which is simply a rearrangement of **equation (5)**.

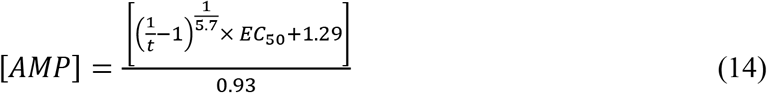

Similarly, we calculated the minimal selective concentration (MSC) as the AMP concentration at 99% fitness (t = 0.99) for wtVIM-2, such that no variant can exhibit more than a 1% increase in fitness over *wt*VIM-2 at this concentration of AMP or lower.

### Nonspecific epistasis of VIM-2 variants

We randomly sampled 200 missense variants, extracted their *EC*_50_ values, and generated all pairwise mutational combinations (_200_C_2_ = 19900 mutant pairs). Since the analysis was focused on the numerical behavior rather than the physical behavior of the enzyme, mutations from the same residue positions were allowed to be combined, though this rarely occurred as only 88 out of 19900 combinations (~0.4%) were from combining the same positions. From these mutant pairs, we used **equation (5)** to calculate the expected fitness effect due to additive changes in fitness relative to *wt*VIM-2. First, the relative change in fitness for each single variant in the pair is calculated according to **equation (15)**, then added together as shown in **equation (16)**.

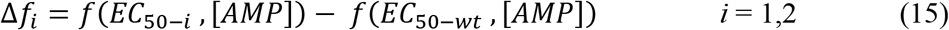

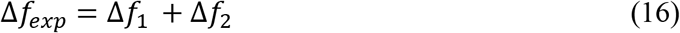

Since the model explicitly defines 0 as the minimum fitness and 1 as the maximum fitness, Δ*f exp* is limited to the maximum change that can occur within this range (−1 to 1).

We also calculate the observed fitness effect according to **equation (5)** as an additive effect in the log2(*EC*_50_) change relative to *wt*VIM-2. First, the relative change in *EC*_50_ is calculated in **equation (17)** and the additive *EC*_50_ effects are used to calculate the new *EC*_50_ in **equation (18)**.

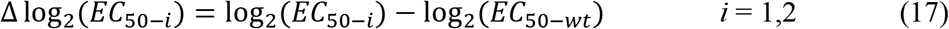

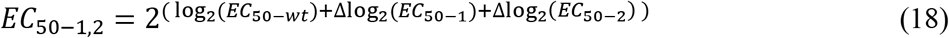

We limit *EC*_50_-1,2 to be no lower than 2.0 (4.0 μg/mL untransformed), as an [AMP] of 2.0 μg/mL is not selective and is not accounted for by **equation (5)**. The observed fitness effect is then calculated according to **equation (19)**.

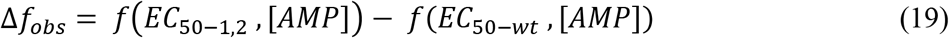

To identify significant epistatic effects, we use the synonymous fitness distribution at each [AMP] as a measure of expected measurement noise. When |Δ*f*obs – Δ*f*exp| < 1.96 × S.D. of the synonymous fitness (after normalization by **equation (13)**), the effect of the mutational pair is considered not significantly epistatic. For mutational pairs with significant epistasis, they are classified as exhibiting positive epistasis when Δ*f*obs > Δ*f*exp, and negative epistasis if Δ*f*obs < Δ*f*exp.

---

**Fig. S1.**
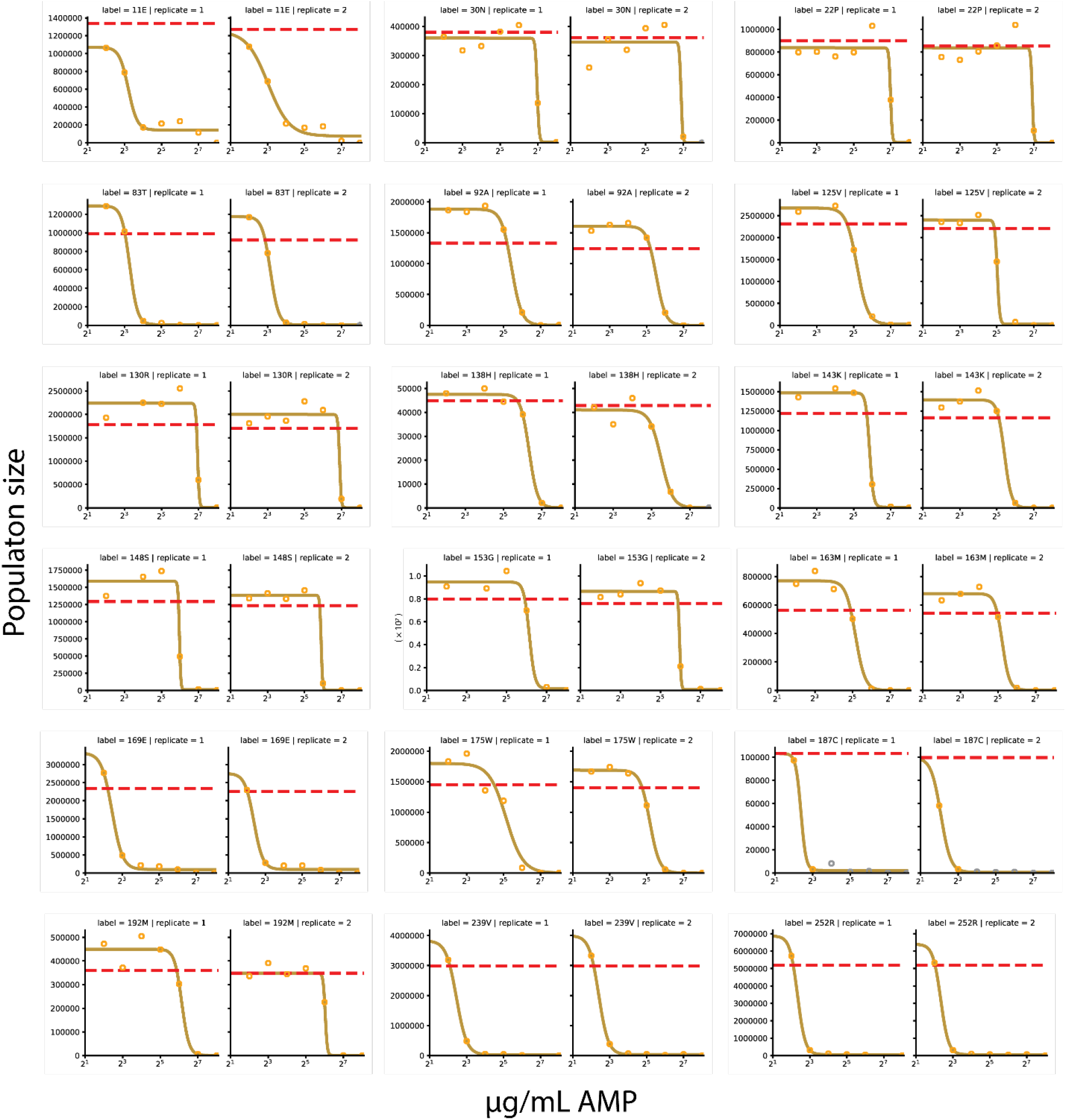
Example dose-response curves for *EC*_50_ calculation based on deep sequencing data. Each pair of plots represent the two replicate dose-response curves of a single variant in the library. The solid curve indicates the results of fitting to the sigmoidal **equation (2)**. The horizontal dashed line marks the average population size of the variant in the non-selected conditions. Dose-response curves shown here were randomly sampled from the entire pool of all variants with a successful fit.

**Fig. S2.**
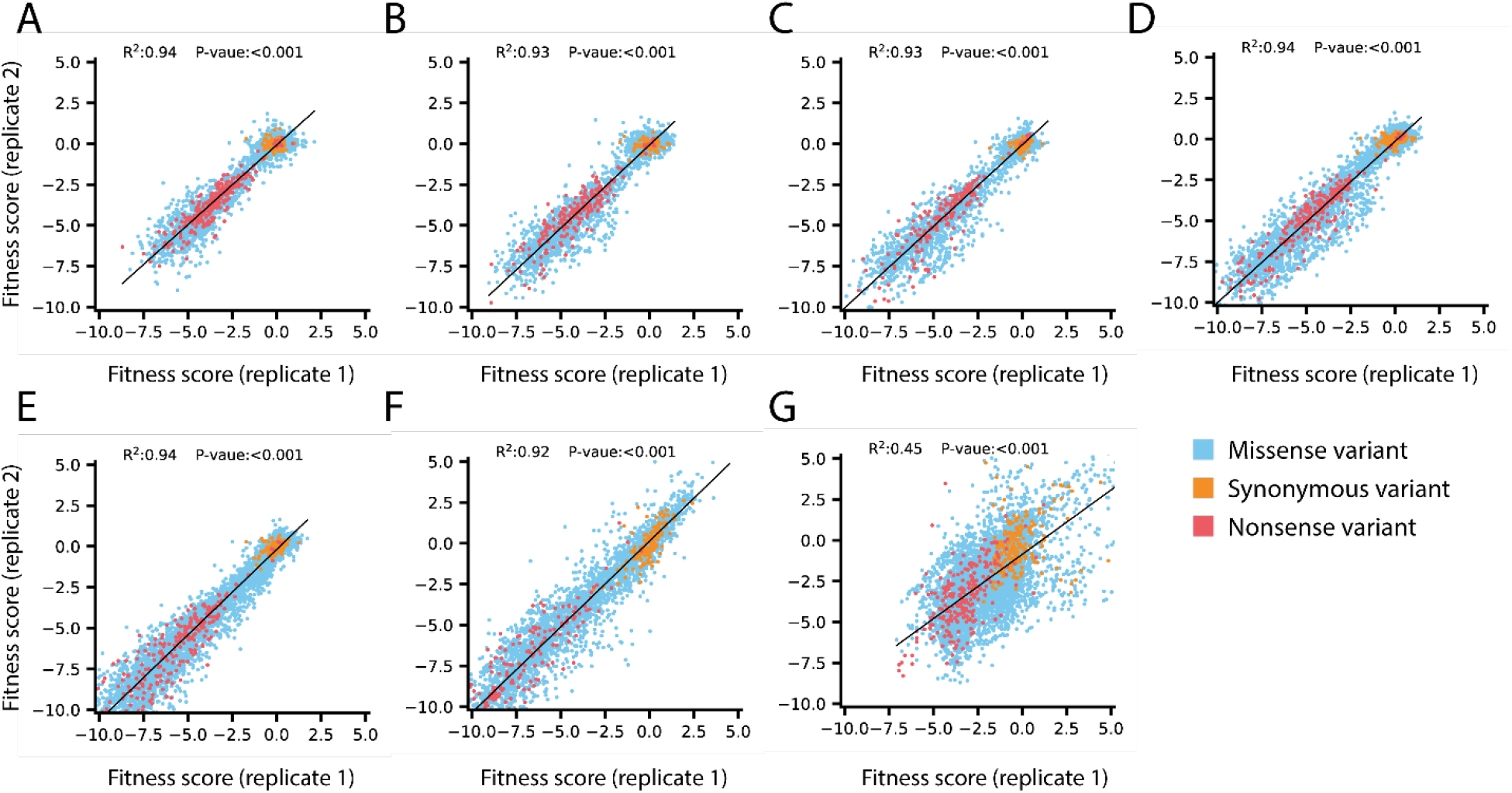
Replicate correlation of DMS fitness scores. Each scatter plot shows the replicate fitness scores of all variants under selection in (**A**) 4.0, (**B**) 8.0, (**C**) 16, (**D**) 32, (**E**) 64, (**F**) 128 and (**G**) 256 μg/mL AMP. In each panel, variants are colored according to mutation type with the legend in the lower right. The solid black line indicate the line of best fit for a linear regression, with the R^2^ and P-value of the regression above each plot.

**Fig. S3.**
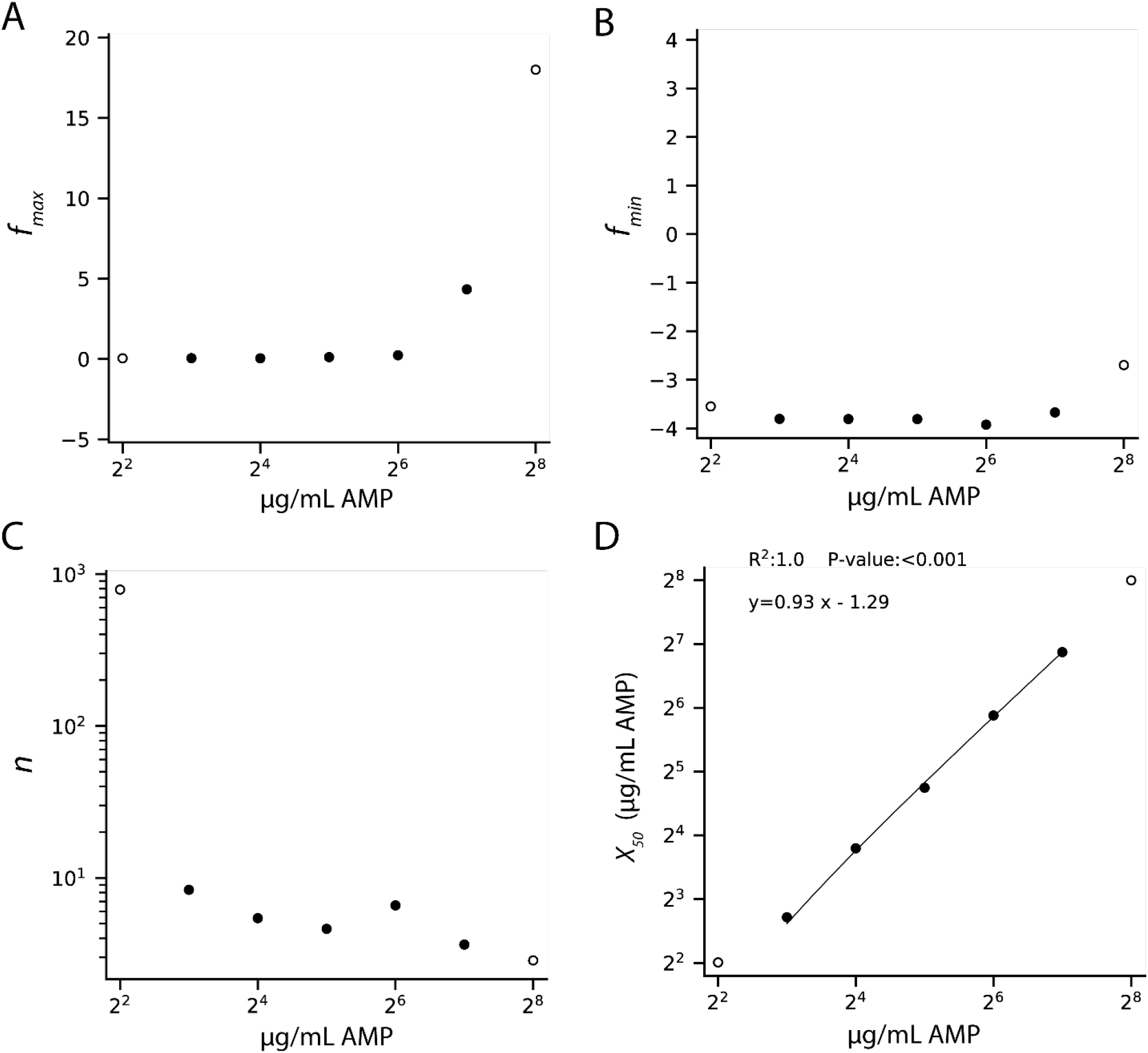
Fit parameters for the phenotype-fitness relationship across AMP concentrations. Each plot shows one of the final fitted value of a parameter in the four-parameter sigmoidal curve in **equation (5)** when fitted to the fitness scores and *EC*_50_ values from the AMP concentration indicated in the x-axis. (**A**) Plot for maximum fitness (*f*_max_) (**B**) Plot for minimum fitness (*f*_min_), (**C**) Plot for Hill coefficient (*n*) (**D**) Plot for the inflection point (*X*_50_). For all plots, solid points represent AMP concentrations where the linear portion of the fit were not limited by the AMP concentration range, while hollow points indicate the opposite. For **D**), a linear regression was conducted with the solid data points. The black line indicates the line of best fit for a linear regression, and the R^2^, P-value and the equation of the fit shown at the top.

**Data S1 (separate file)**

Fitness scores of VIM-2 variants across AMP concentrations.

**Data S2 (separate file)**

EC50 of all VIM-2 variants.

**Data S3 (separate file)**

EC50 of VIM-2 variants measured individually.

## References and Notes

1. A. C. Dalziel, S. M. Rogers, P. M. Schulte, Linking genotypes to phenotypes and fitness: how mechanistic biology can inform molecular ecology. Mol. Ecol. 18, 4997–5017 (2009).

2. M. Kaltenbach, N. Tokuriki, Dynamics and constraints of enzyme evolution. J. Exp. Zool. B Mol. Dev. Evol. 322, 468–487 (2014).

3. X. Yi, A. M. Dean, Adaptive Landscapes in the Age of Synthetic Biology. Mol. Biol. Evol. 36, 890–907 (2019).

4. H. Kemble, P. Nghe, O. Tenaillon, Recent insights into the genotype–phenotype relationship from massively parallel genetic assays. Evolutionary Applications. 12, 1721–1742 (2019).

5. S. Bershtein, A. W. Serohijos, E. I. Shakhnovich, Bridging the physical scales in evolutionary biology: from protein sequence space to fitness of organisms and populations. Curr. Opin. Struct. Biol. 42, 31–40 (2017).

6. M. A. DePristo, D. M. Weinreich, D. L. Hartl, Missense meanderings in sequence space: a biophysical view of protein evolution. Nature Reviews Genetics. 6, 678 (2005).

7. C. Pál, B. Papp, M. J. Lercher, An integrated view of protein evolution. Nat. Rev. Genet. 7, 337–348 (2006).

8. J. V. Rodrigues, S. Bershtein, A. Li, E. R. Lozovsky, D. L. Hartl, E. I. Shakhnovich, Biophysical principles predict fitness landscapes of drug resistance. Proc Natl Acad Sci U S A. 113, E1470–1478 (2016).

9. D. L. Hartl, D. E. Dykhuizen, A. M. Dean, Limits of adaptation: the evolution of selective neutrality. Genetics. 111, 655–674 (1985).

10. M. Lunzer, S. P. Miller, R. Felsheim, A. M. Dean, The Biochemical Architecture of an Ancient Adaptive Landscape. Science. 310, 499–501 (2005).

11. E. Lundin, P.-C. Tang, L. Guy, J. Näsvall, D. I. Andersson, Experimental Determination and Prediction of the Fitness Effects of Random Point Mutations in the Biosynthetic Enzyme HisA. Mol Biol Evol. 35, 704–718 (2018).

12. N. Tokuriki, D. S. Tawfik, Stability effects of mutations and protein evolvability. Current Opinion in Structural Biology. 19, 596–604 (2009).

13. K. S. Sarkisyan, D. A. Bolotin, M. V. Meer, D. R. Usmanova, A. S. Mishin, G. V. Sharonov, D. N. Ivankov, N. G. Bozhanova, M. S. Baranov, O. Soylemez, N. S. Bogatyreva, P. K. Vlasov, E. S. Egorov, M. D. Logacheva, A. S. Kondrashov, D. M. Chudakov, E. V. Putintseva, I. Z. Mamedov, D. S. Tawfik, K. A. Lukyanov, F. A. Kondrashov, Local fitness landscape of the green fluorescent protein. Nature. 533, 397–401 (2016).

14. P. Dasmeh, A. W. R. Serohijos, Estimating the contribution of folding stability to nonspecific epistasis in protein evolution. Proteins: Structure, Function, and Bioinformatics. 86, 1242–1250 (2018).

15. S. Bershtein, M. Segal, R. Bekerman, N. Tokuriki, D. S. Tawfik, Robustness–epistasis link shapes the fitness landscape of a randomly drifting protein. Nature. 444, 929–932 (2006).

16. V. O. Pokusaeva, D. R. Usmanova, E. V. Putintseva, L. Espinar, K. S. Sarkisyan, A. S. Mishin, N. S. Bogatyreva, D. N. Ivankov, A. V. Akopyan, S. Y. Avvakumov, I. S. Povolotskaya, G. J. Filion, L. B. Carey, F. A. Kondrashov, An experimental assay of the interactions of amino acids from orthologous sequences shaping a complex fitness landscape. PLOS Genetics. 15, e1008079 (2019).

17. G. Diss, B. Lehner, The genetic landscape of a physical interaction. eLife. 7, e32472 (2018).

18. C. S. Wylie, E. I. Shakhnovich, A biophysical protein folding model accounts for most mutational fitness effects in viruses. Proc Natl Acad Sci U S A. 108, 9916–9921 (2011).

19. S. Bershtein, W. Mu, A. W. R. Serohijos, J. Zhou, E. I. Shakhnovich, Protein Quality Control Acts on Folding Intermediates to Shape the Effects of Mutations on Organismal Fitness. Molecular Cell. 49, 133–144 (2013).

20. M. A. Stiffler, D. R. Hekstra, R. Ranganathan, Evolvability as a Function of Purifying Selection in TEM-1 β-Lactamase. Cell. 160, 882–892 (2015).

21. J. D. Mehlhoff, F. W. Stearns, D. Rohm, B. Wang, E.-Y. Tsou, N. Dutta, M.-H. Hsiao, C. E. Gonzalez, A. F. Rubin, M. Ostermeier, Collateral fitness effects of mutations. PNAS. 117, 11597–11607 (2020).

22. A. M. Dean, A molecular investigation of genotype by environment interactions. Genetics. 139, 19–33 (1995).

23. D. M. Fowler, C. L. Araya, S. J. Fleishman, E. H. Kellogg, J. J. Stephany, D. Baker, S. Fields, High-resolution mapping of protein sequence-function relationships. Nature Methods. 7, 741–746 (2010).

24. D. M. Fowler, J. J. Stephany, S. Fields, Measuring the activity of protein variants on a large scale using deep mutational scanning. Nat Protoc. 9, 2267–2284 (2014).

25. A. Melnikov, P. Rogov, L. Wang, A. Gnirke, T. S. Mikkelsen, Comprehensive mutational scanning of a kinase *in vivo* reveals substrate-dependent fitness landscapes. Nucleic Acids Research. 42, e112–e112 (2014).

26. E. E. Wrenbeck, L. R. Azouz, T. A. Whitehead, Single-mutation fitness landscapes for an enzyme on multiple substrates reveal specificity is globally encoded. Nature Communications. 8, 15695 (2017).

27. D. Mavor, K. Barlow, S. Thompson, B. A. Barad, A. R. Bonny, C. L. Cario, G. Gaskins, Z. Liu, L. Deming, S. D. Axen, E. Caceres, W. Chen, A. Cuesta, R. E. Gate, E. M. Green, K. R. Hulce, W. Ji, L. R. Kenner, B. Mensa, L. S. Morinishi, S. M. Moss, M. Mravic, R. K. Muir, S. Niekamp, C. I. Nnadi, E. Palovcak, E. M. Poss, T. D. Ross, E. C. Salcedo, S. K. See, M. Subramaniam, A. W. Wong, J. Li, K. S. Thorn, S. Ó. Conchúir, B. P. Roscoe, E. D. Chow, J. L. DeRisi, T. Kortemme, D. N. Bolon, J. S. Fraser, Determination of ubiquitin fitness landscapes under different chemical stresses in a classroom setting. Elife. 5 (2016), doi:10.7554/eLife.15802.

28. L. Noda-García, D. Davidi, E. Korenblum, A. Elazar, E. Putintseva, A. Aharoni, D. S. Tawfik, Chance and pleiotropy dominate genetic diversity in complex bacterial environments. Nat Microbiol. 4, 1221–1230 (2019).

29. T. N. Starr, L. K. Picton, J. W. Thornton, Alternative evolutionary histories in the sequence space of an ancient protein. Nature. 549, 409–413 (2017).

30. T. N. Starr, A. J. Greaney, S. K. Hilton, D. Ellis, K. H. D. Crawford, A. S. Dingens, M. J. Navarro, J. E. Bowen, M. A. Tortorici, A. C. Walls, N. P. King, D. Veesler, J. D. Bloom, Deep Mutational Scanning of SARS-CoV-2 Receptor Binding Domain Reveals Constraints on Folding and ACE2 Binding. Cell. 182, 1295–1310.e20 (2020).

31. R. T. Hietpas, J. D. Jensen, D. N. A. Bolon, Experimental illumination of a fitness landscape. Proc. Natl. Acad. Sci. U.S.A. 108, 7896–7901 (2011).

32. E. Firnberg, J. W. Labonte, J. J. Gray, M. Ostermeier, A Comprehensive, High-Resolution Map of a Gene’s Fitness Landscape. Molecular Biology and Evolution. 31, 1581–1592 (2014).

33. H. Jacquier, A. Birgy, H. L. Nagard, Y. Mechulam, E. Schmitt, J. Glodt, B. Bercot, E. Petit, J. Poulain, G. Barnaud, P.-A. Gros, O. Tenaillon, Capturing the mutational landscape of the beta-lactamase TEM-1. PNAS. 110, 13067–13072 (2013).

34. J. R. Klesmith, J.-P. Bacik, E. E. Wrenbeck, R. Michalczyk, T. A. Whitehead, Trade-offs between enzyme fitness and solubility illuminated by deep mutational scanning. Proceedings of the National Academy of Sciences. 114, 2265–2270 (2017).

35. J. Z. Chen, D. M. Fowler, N. Tokuriki, Comprehensive exploration of the translocation, stability and substrate recognition requirements in VIM-2 lactamase. eLife. 9, e56707 (2020).

36. J.-Y. van der Meer, H. Poddar, B.-J. Baas, Y. Miao, M. Rahimi, A. Kunzendorf, R. van Merkerk, P. G. Tepper, E. M. Geertsema, A.-M. W. H. Thunnissen, W. J. Quax, G. J. Poelarends, Using mutability landscapes of a promiscuous tautomerase to guide the engineering of enantioselective Michaelases. Nat Commun. 7, 1–16 (2016).

37. S. Thompson, Y. Zhang, C. Ingle, K. A. Reynolds, T. Kortemme, Altered expression of a quality control protease in E. coli reshapes the in vivo mutational landscape of a model enzyme. eLife. 9, e53476 (2020).

38. H. Kemble, C. Eisenhauer, A. Couce, A. Chapron, M. Magnan, G. Gautier, H. L. Nagard, P. Nghe, O. Tenaillon, Flux, toxicity, and expression costs generate complex genetic interactions in a metabolic pathway. Science Advances. 6, eabb2236 (2020).

39. L. M. Starita, J. N. Pruneda, R. S. Lo, D. M. Fowler, H. J. Kim, J. B. Hiatt, J. Shendure, P. S. Brzovic, S. Fields, R. E. Klevit, Activity-enhancing mutations in an E3 ubiquitin ligase identified by high-throughput mutagenesis. Proceedings of the National Academy of Sciences. 110, E1263–E1272 (2013).

40. J. M. Flynn, A. Rossouw, P. Cote-Hammarlof, I. Fragata, D. Mavor, C. Hollins III, C. Bank, D. N. Bolon, Comprehensive fitness maps of Hsp90 show widespread environmental dependence. eLife. 9, e53810 (2020).

41. A. Horovitz, R. C. Fleisher, T. Mondal, Double-mutant cycles: new directions and applications. Current Opinion in Structural Biology. 58, 10–17 (2019).

42. V. H. Salinas, R. Ranganathan, Coevolution-based inference of amino acid interactions underlying protein function. eLife. 7, e34300 (2018).

43. D. Mavor, K. A. Barlow, D. Asarnow, Y. Birman, D. Britain, W. Chen, E. M. Green, L. R. Kenner, B. Mensa, L. S. Morinishi, C. A. Nelson, E. M. Poss, P. Suresh, R. Tian, T. Arhar, B. E. Ary, D. P. Bauer, I. D. Bergman, R. M. Brunetti, C. M. Chio, S. A. Dai, M. S. Dickinson, S. K. Elledge, C. V. M. Helsell, N. L. Hendel, E. Kang, N. Kern, M. S. Khoroshkin, L. L. Kirkemo, G. R. Lewis, K. Lou, W. M. Marin, A. M. Maxwell, P. F. McTigue, D. Myers-Turnbull, T. L. Nagy, A. M. Natale, K. Oltion, S. Pourmal, G. K. Reder, N. J. Rettko, P. J. Rohweder, D. M. C. Schwarz, S. K. Tan, P. V. Thomas, R. W. Tibble, J. P. Town, M. K. Tsai, F. S. Ugur, D. R. Wassarman, A. M. Wolff, T. S. Wu, D. Bogdanoff, J. Li, K. S. Thorn, S. O’Conchúir, D. L. Swaney, E. D. Chow, H. D. Madhani, S. Redding, D. N. Bolon, T. Kortemme, J. L. DeRisi, M. Kampmann, J. S. Fraser, Extending chemical perturbations of the ubiquitin fitness landscape in a classroom setting reveals new constraints on sequence tolerance. Biol Open. 7 (2018), doi:10.1242/bio.036103.

44. L. Rockah-Shmuel, Á. Tóth-Petróczy, D. S. Tawfik, Systematic Mapping of Protein Mutational Space by Prolonged Drift Reveals the Deleterious Effects of Seemingly Neutral Mutations. PLoS Comput. Biol. 11, e1004421 (2015).

45. V. E. Gray, R. J. Hause, J. Luebeck, J. Shendure, D. M. Fowler, Quantitative Missense Variant Effect Prediction Using Large-Scale Mutagenesis Data. Cell Syst. 6, 116–124.e3 (2018).

46. E. Gullberg, S. Cao, O. G. Berg, C. Ilbäck, L. Sandegren, D. Hughes, D. I. Andersson, Selection of resistant bacteria at very low antibiotic concentrations. PLoS Pathog. 7, e1002158 (2011).

47. S. Westhoff, T. M. van Leeuwe, O. Qachach, Z. Zhang, G. P. van Wezel, D. E. Rozen, The evolution of no-cost resistance at sub-MIC concentrations of streptomycin in Streptomyces coelicolor. ISME J. 11, 1168–1178 (2017).

48. A. Liu, A. Fong, E. Becket, J. Yuan, C. Tamae, L. Medrano, M. Maiz, C. Wahba, C. Lee, K. Lee, K. P. Tran, H. Yang, R. M. Hoffman, A. Salih, J. H. Miller, Selective advantage of resistant strains at trace levels of antibiotics: a simple and ultrasensitive color test for detection of antibiotics and genotoxic agents. Antimicrob. Agents Chemother. 55, 1204–1210 (2011).

49. W. Jasinska, M. Manhart, J. Lerner, L. Gauthier, A. W. R. Serohijos, S. Bershtein, Chromosomal barcoding of E. coli populations reveals lineage diversity dynamics at high resolution. Nature Ecology & Evolution. 4, 437–452 (2020).

50. C. Fröhlich, J. A. Gama, K. Harms, V. H. A. Hirvonen, B. A. Lund, M. W. van der Kamp, P. J. Johnsen, Ø. Samuelsen, H.-K. S. Leiros, bioRxiv, in press, doi:10.1101/2020.12.01.404343.

51. D. I. Andersson, D. Hughes, Microbiological effects of sublethal levels of antibiotics. Nature Reviews Microbiology. 12, 465–478 (2014).

52. F. Baquero, M. C. Negri, M. I. Morosini, J. Blázquez, Selection of very small differences in bacterial evolution. Int Microbiol. 1, 295–300 (1998).

53. K. Drlica, X. Zhao, Mutant Selection Window Hypothesis Updated. Clin Infect Dis. 44, 681–688 (2007).

54. J. M. Blondeau, New concepts in antimicrobial susceptibility testing: the mutant prevention concentration and mutant selection window approach. Vet Dermatol. 20, 383–396 (2009).

55. S. G. Das, S. O. Direito, B. Waclaw, R. J. Allen, J. Krug, Predictable properties of fitness landscapes induced by adaptational tradeoffs. eLife. 9, e55155 (2020).

56. T. N. Starr, J. W. Thornton, Epistasis in protein evolution. Protein Sci. 25, 1204–1218 (2016).

57. J. Domingo, P. Baeza-Centurion, B. Lehner, The Causes and Consequences of Genetic Interactions (Epistasis). Annu Rev Genomics Hum Genet. 20, 433–460 (2019).

58. G. Yang, D. W. Anderson, F. Baier, E. Dohmen, N. Hong, P. D. Carr, S. C. L. Kamerlin, C. J. Jackson, E. Bornberg-Bauer, N. Tokuriki, Higher-order epistasis shapes the fitness landscape of a xenobiotic-degrading enzyme. Nature Chemical Biology. 15, 1120–1128 (2019).

59. F. Baier, N. Hong, G. Yang, A. Pabis, C. M. Miton, A. Barrozo, P. D. Carr, S. C. Kamerlin, C. J. Jackson, N. Tokuriki, Cryptic genetic variation shapes the adaptive evolutionary potential of enzymes. Elife. 8 (2019), doi:10.7554/eLife.40789.

60. C. Bank, R. T. Hietpas, J. D. Jensen, D. N. A. Bolon, A systematic survey of an intragenic epistatic landscape. Mol Biol Evol. 32, 229–238 (2015).

61. C. L. Araya, D. M. Fowler, W. Chen, I. Muniez, J. W. Kelly, S. Fields, A fundamental protein property, thermodynamic stability, revealed solely from large-scale measurements of protein function. Proceedings of the National Academy of Sciences. 109, 16858–16863 (2012).

62. C. A. Olson, N. C. Wu, R. Sun, A comprehensive biophysical description of pairwise epistasis throughout an entire protein domain. Curr Biol. 24, 2643–2651 (2014).

63. A. C. Palmer, E. Toprak, M. Baym, S. Kim, A. Veres, S. Bershtein, R. Kishony, Delayed commitment to evolutionary fate in antibiotic resistance fitness landscapes. Nat Commun. 6, 7385 (2015).

64. C. M. Miton, N. Tokuriki, How mutational epistasis impairs predictability in protein evolution and design. Protein Sci. 25, 1260–1272 (2016).

65. B. Steinberg, M. Ostermeier, Environmental changes bridge evolutionary valleys. Sci. Adv. 2, e1500921 (2016).

66. M. P. Zwart, M. F. Schenk, S. Hwang, B. Koopmanschap, N. de Lange, L. van de Pol, T. T. T. Nga, I. G. Szendro, J. Krug, J. A. G. M. de Visser, Unraveling the causes of adaptive benefits of synonymous mutations in TEM-1 β-lactamase. Heredity. 121, 406–421 (2018).

67. R. C. Edgar, H. Flyvbjerg, Error filtering, pair assembly and error correction for next-generation sequencing reads. Bioinformatics. 31, 3476–3482 (2015).

68. J. L. Sebaugh, P. D. McCray, Defining the linear portion of a sigmoid-shaped curve: bend points. Pharmaceutical Statistics. 2, 167–174 (2003).

